# Concomitant AD and DLB pathologies shape subfield microglia responses in the hippocampus

**DOI:** 10.1101/2022.01.06.475218

**Authors:** Sonja Fixemer, Corrado Ameli, Gaël Hammer, Luis Salamanca, Oihane Uriarte Huarte, Chantal Schwartz, Jean-Jacques Gérardy, Naguib Mechawar, Alexander Skupin, Michel Mittelbronn, David Bouvier

**Affiliations:** Luxembourg Centre for Systems Biomedicine (LCSB), University of Luxembourg, Belval, Luxembourg; Luxembourg Center of Neuropathology (LCNP), Dudelange, Luxembourg; Laboratoire national de santé (LNS), National Center of Pathology (NCP), Dudelange, Luxembourg; Swiss Data Science Center, ETH Zürich, Universitätstrasse 25, Zürich, Switzerland; Douglas Mental Health University Institute, Department of Psychiatry, McGill University, Montreal, Quebec, Canada; Centre for Research in Neuroscience, Department of Neurology and Neurosurgery, The Research Institute of the McGill University Health Centre, Montreal General Hospital, Montreal, Quebec, Canada; Department of Oncology (DONC), Luxembourg Institute of Health (LIH), Luxembourg; Department of Life Sciences and Medicine (DLSM), University of Luxembourg, Esch-sur-Alzette, Luxembourg; Faculty of Science, Technology and Medicine (FSTM), University of Luxembourg, Esch-sur-Alzette, Luxembourg

**Keywords:** Alzheimer’s Disease, Dementia with Lewy Bodies, Hippocampus, Microglia, Amyloid-beta, Tau, Alpha-synuclein

## Abstract

Hippocampal alteration is at the centre of memory decline in the most common age-related neurodegenerative diseases: Alzheimer’s disease (AD) and Dementia with Lewy Bodies (DLB). However, the subregional deterioration of the hippocampus differs between both diseases with more severe atrophy in the CA1 subfield of the AD patients. How AD and DLB-typical pathologies compose the various local microenvironment of the hippocampus across AD and DLB needs to be further explored to understand this process. Additionally, microglia responses could further impact the atrophy rate. Some studies suggest that microglia react differently according to the underlying neurodegenerative disorder. How microglia are transformed across hippocampal subfields in AD and DLB, and how their changes are associated with disease-typical pathologies remains to be determined. To these purposes, we performed a volumetric analysis of phospho-Tau (P-Tau), Amyloid-ß (Aß), and phospho-α-Synuclein (P-Syn) loads, quantified and classified microglia according to distinct morphological phenotypes using high-resolution confocal 3D microscopy of hippocampal CA1, CA3 and DG/CA4 subfields of late-onset AD (n=10) and DLB (n=8) as well as age-matched control samples (n=11).

We found that each of the Tau, Aß and Synuclein pathologies followed a specific subregional distribution, relatively preserved across AD and DLB. P-Tau, Aß and P-Syn burdens were significantly exacerbated in AD, with Tau pathology being particularly severe in the AD CA1. P-Tau and P-Syn burdens were highly correlated across subfields and conditions (R2Spear = 0.79; P < 0.001) and result from a local co-distribution of P-Tau and P-Syn inclusions in neighbouring neurons, with only a low proportion of double-positive cells.

In parallel, we assessed the changes of the microglia responses by measuring 16 morphological features of more than 35,000 individual microglial cells and classifying them into seven-distinct morphological clusters. We found microglia features- and clusters-variations subfield- and condition-dependent. Two of the seven morphological clusters, with more amoeboid and less branched forms, were identified as disease-enriched and found to be further increased in AD. Interestingly, some microglial features or clusters were associated with one but more often with a combination of two pathologies in a subfield-dependent manner.

In conclusion, our study shows a multimodal association of the hippocampal microglia responses with the co-occurrence, distribution and severity of AD and DLB pathologies. In DLB hippocampi, pathological imprint and microglia responses follow AD trends but with lesser severity. Our study suggests that the increased pathological burdens of P-Tau and P-Syn and associated microglia alterations are involved in a more severe deterioration of the CA1 in AD as compared to DLB.

## Introduction

The decline of memory is an overlapping symptom in age-related neurodegenerative diseases (NDDs)^1^. The hippocampus is at the centre of numerous cognitive processes and is essential for spatial and episodic memory formation and consolidation^2–4^. It is one of the most severely affected brain regions in Alzheimer’s disease (AD) and Dementia with Lewy Bodies (DLB)^5– 7^ but the causes of its vulnerability are still poorly understood. The loss of hippocampal volume is associated with memory impairment in normal ageing^8^ and is predictive of AD and associated dementia^9^. Its architecture has been widely studied for its tri-synaptic loop circuitry and is composed of 5 subfields, the *dentate gyrus* (DG) as well as the *Cornu ammonis* (CA) fields CA1, CA2, CA3 and CA4^10^. Hippocampal subfields are distinct in their function and cellular composition and could be dedicated to or specialized in distinct types or sequences of memory processes^11–13^. Interestingly, the atrophy of the hippocampus follows specific subregional patterns between AD and DLB cases, with the CA1 subfield particularly affected by neuronal loss in AD^14–19^. Understanding the mechanisms that lead to subregional hippocampal vulnerability and deterioration in AD and DLB could unveil personalized therapeutic strategies to alleviate the progression of dementia in NDDs.

At the cellular level many factors could be involved in the accelerated deterioration of CA1 in AD. NDDs are associated with extra or intra-cellular deposits of misfolded proteins across the CNS parenchyma that are correlated with disease progression. Extracellular senile plaques made of Amyloid-ß (Aß) peptides, intracellular neurofibrillary tangles (NFT) built from hyperphosphorylated Tau (P-Tau) and intracellular aggregation of phosphorylated α-Synuclein (P-Syn) such as Lewy bodies^20^, are diagnostic neuropathological hallmarks in NDDs ^21–24^. The overlapping of Tau, Aß and Synuclein pathologies is frequent in DLB but also in AD in approximately two-thirds of the cases^25–32^. It raises numerous questions about how intertwined neurodegenerative mechanisms are across conditions and their impact on the progression of the disease. Microglia, the innate immune cells of the CNS, regulate homeostasis by clearing pathogens, cell debris and dying cells in the brain parenchyma ^33^. In cell cultures and rodent models, Tau, Aß and Synuclein pathologies are all three strongly associated with microglial activation and responses ^34–40^. If these interactions prime microglial cells to alleviate or exacerbate NDDs progression is still under discussion. Microglia aggregation is typically observed around Aß plaques, where microglia have been involved in Aß 1-42 peptide and oligomer clearance but also in the formation and maintenance of Aß plaques^41–45^. Microglia attached to Aß plaques also display distinct molecular signatures ^40,46,47^. Their relationship with Tau pathology spreading is still under debate with contradictive results emerging from studies using different mouse models^48–50^. Furthermore, recent studies report a potential role of microglia in P-Syn clearance ^51^ and its cell-to-cell transfer^52^. Additionally, the identification by genome-wide association studies (GWAS) of risk loci associated with genes involved in their physiology and responses such as CR1 (coding for the Complement receptor type 1), SPI1 (Transcription factor PU.1), TREM2 (Triggering receptor expressed on myeloid cells 2) and CD33^53^emphasizes on the association of their functions with AD onset. Maladaptive responses such as uncontrolled phagocytic activities or oversecretion of pro-inflammatory factors such as the tumor necrosis factor-α (TNF-α), interleukin-1β (IL-1 β), and reactive oxygen species (ROS), could accelerate neuronal deterioration^42^. Furthermore, reports of regional and/or temporal heterogeneity of microglia signatures across mouse models and neurodegenerative conditions in human increase the complexity to define how and when microglia play beneficial or detrimental roles^54–57^ and raise the importance to better define their changes in association with the mixed pathologies contexts of patient samples. Some studies have already highlighted differences in changes and activation in the microglia network between AD and DLB^58–63^ with the microglia in DLB samples seemingly less altered and responsive. However, how microglia responses are associated to AD- and DLB-pathology burdens in the human hippocampus is still an open question.

In this study, we investigated how the co-occurrence of Tau, Aß and Synuclein pathologies modifies the subregional pathological burden and microglia morphological changes across the hippocampus of AD and DLB cases. We immunostained thick sections with P-Tau, Aß, P-Syn or Iba1 antibodies and employed high-content 3D confocal microscopy combined with image analysis tools such as our Microglia and Immune Cells Morphologies Analyser and Classifier (MIC-MAC) pipeline on a collection of neuropathologically diagnosed AD and DLB cases, and age-matched controls (CTLs) samples. Our analysis revealed a specific distribution pattern across hippocampal subfields for each pathology, a strong correlation between P-Tau and P-Syn loads, as well as a fine-tuned association between microglia morphological changes and co-occurrence and severity of AD- and DLB-pathologies across subregions and conditions.

## Materials and methods

### Human brain samples and processing

All experiments involving human tissues were conducted in accordance with the guidelines approved by the Ethics Board of the Douglas-Bell Brain Bank (Douglas Mental Health University Institute, Montréal, QC, Canada) and the Ethic Panel of the University of Luxembourg (ERP 16-037 and 21-009). All anonymized autopsy brain samples were obtained from the Douglas-Bell Brain Bank (see Table 1). Hippocampal samples (median part of the hippocampus, between the uncus and *corpus geniculatum laterale*) were dissected from cases of neuropathologically confirmed AD and DLB, as well as from age matched CTLs. Samples were evaluated regarding Aß plaques, NFT and/or alpha-synuclein and assessed according to Braak and ABC staging ^21,64^ or Mc Keith staging criteria ^65^ by a neuropathologist affiliated to the Douglas-Bell Brain Bank. This study comprises age-matched CTLs with no history of dementia. However, some CTL cases showed low level of neuropathological abnormalities such as AD pathology. Human brain samples from the Douglas-Bell brain bank used for the volumetric study were preserved in 10% formalin until processing as indicated in Table 1.

**Table 1.**
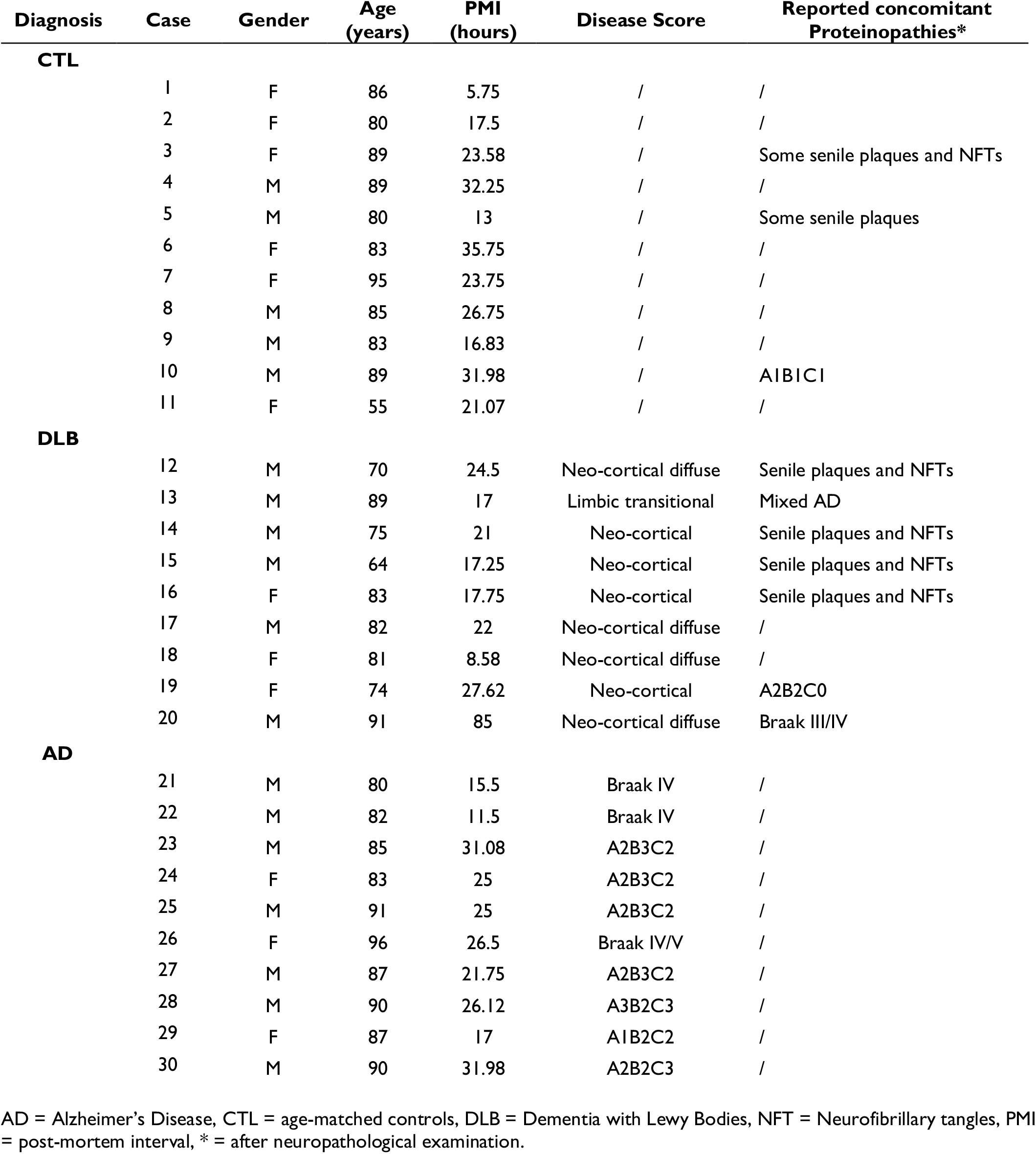
Demographic and neuropathological data of human samples.

### Post-mortem case description

The description of the samples is detailed in Table 1. We ran our image-based analysis on 11 age-matched CTLs, with no or low levels of neuropathological alterations as well as 10 AD (A1B2C2 to severe stages A2B3C2 or Braak IV) and 8 DLB (limbic transitional to neocortical diffuse) cases bearing often a certain amount of AD pathologies. The average year of death and post-mortem delay (PMD) did not differ significantly between groups and are 83.1 years and 22.56 hours for CTLs; 87.1 and 23.14 for AD; and 77.3 and 19.4 for DLB. Gender representation is slightly unbalanced with female cases being respectively 54.5% for CTLs, 30% for AD and 37.5% for DLB. Average time spent in fixative was also approximatively similar for specimens of all groups (CTLs, 14.7 years; AD, 11.1 and DLB, 12.25).

### Immunohistochemistry

One sample from the Luxembourg Brain Bank was used to validate P-Syn antibodies on 3 µm Formalin-fixed paraffin-embedded (FFPE) sections. Briefly, sections were processed by an automated immunostainer (Omnis Immunostainer, Agilent, Glostrup, Denmark) with primary antibodies against PS129-synuclein. Heat retrieval was performed with the Envision Flex Tris low pH buffer (Agilent) 30 min at 97°C on board, primary antibodies were incubated 20 min at RT, and detection was performed with the Envision Flex Detection kit (Agilent) based on the DAB/HRP substrate system.

Before sectioning, human hippocampal brain samples from the Douglas-Bell Brain Bank preserved in fixative were washed in Phosphate Buffer Saline (PBS) and cryo-preserved in 30% sucrose in PBS for 36 h approximately. Human samples were embedded in M-1 embedding matrix (Thermo scientific, USA) and cut into 50 µm to 100 µm thick slices on a sliding freezing microtome (LeicaSM2010R) and preserved at -20°C in a cryoprotectant solution (Ethylene glycol (30%), and glycerol (30%) in 0.05 M phosphate buffer (PB, pH 7.4)) until processed for immunohistochemistry experiments. Immunostainings were performed as previously described ^66,67^. In brief, after overnight UV irradiation (UV lamp Ushio, 30 Watt), sections were permeabilized for 30 min with 0.5% Triton-X 100 in PBS 1X. Subsequently, free-floating sections were incubated for 2 h with blocking solution (0.5% Triton-X 100 and 2% horse serum in PBS 1X) at room temperature (RT), incubated with primary antibodies in blocking solution for 72 h at 4°C. Sections were then washed three times for 10 min in PBS 1X, and subsequently incubated in 0.5% triton-X 100/PBS 1X at RT for 2 h with fluorophore coupled secondary antibodies. Some slices were additionally incubated for 20 min with the fluorescent DNA dye DRAQ7™ (1:100, Cell Signaling #7406) at RT and washed twice for 10 min in 0.1 M PB (pH 7.4). Sections were finally washed twice for 10 min in 0.1 M PB (pH 7.4) prior to mounting on glass slides using ProLong Gold Antifade reagent (Invitrogen). All references for primary and secondary antibodies can be found in Table 2, respectively Table 3.

**Table 2.**
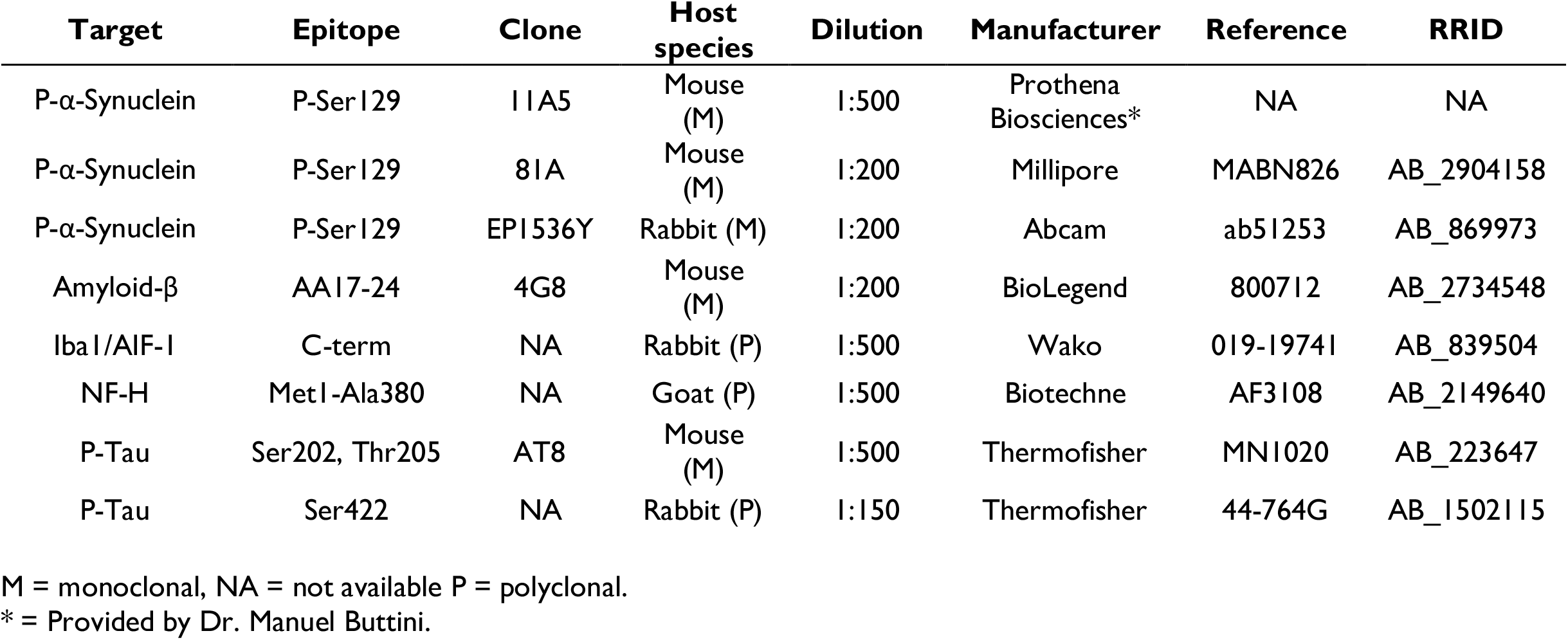
Primary antibodies.

**Table 3.**
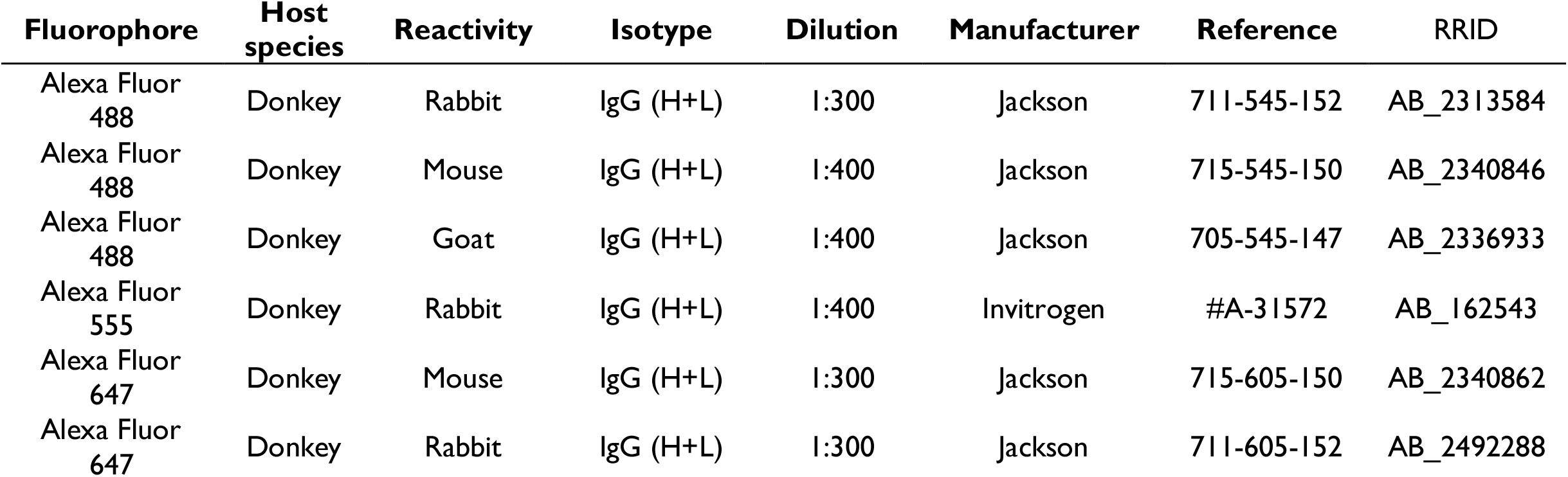
Secondary antibodies.

| Fluorophore | Host species | Reactivity | Isotype | Dilution | Manufacturer | Reference | RRID |
| --- | --- | --- | --- | --- | --- | --- | --- |
| Alexa Fluor 488 | Donkey | Rabbit | IgG (H+L) | 1:300 | Jackson | 711-545-152 | AB_2313584 |
| Alexa Fluor 488 | Donkey | Mouse | IgG (H+L) | 1:400 | Jackson | 715-545-150 | AB_2340846 |
| Alexa Fluor 488 | Donkey | Goat | IgG (H+L) | 1:400 | Jackson | 705-545-147 | AB_2336933 |
| Alexa Fluor 555 | Donkey | Rabbit | IgG (H+L) | 1:400 | Invitrogen | #A-31572 | AB_162543 |
| Alexa Fluor 647 | Donkey | Mouse | IgG (H+L) | 1:300 | Jackson | 715-605-150 | AB_2340862 |
| Alexa Fluor 647 | Donkey | Rabbit | IgG (H+L) | 1:300 | Jackson | 711-605-152 | AB_2492288 |

### Image acquisition and analysis

#### 3D image acquisition

Confocal images were captured on a Zeiss LSM 710 or LSM 800 confocal system with a 20x air objective. 3D tile scans (z-steps of 1µm) of human hippocampal subfields from age-matched CTLs, DLB and AD patients were stitched via Zeiss or Imaris Stitcher and visualized with Imaris 9.5 and 9.6 (Oxford Instrument). 3D stacks of Iba1 for microglia morphology and for P-Tau, Aß and P-Syn inclusions were acquired on serial sections in the anatomically defined regions of interest representative of the entire layered structure of the CA1, CA3 and DG/CA4: from *Stratum oriens* to *Stratum moleculare* for CA1 and CA3, and of the DG including the hilus (CA4) as indicated in Supplementary Fig. 1.

#### Volumetric quantification of P-Tau, Aß and P-Syn pathologies

To improve homogeneity in the analysis, for each marker, samples were stained following the same combination of primary and secondary antibodies (respectively AT8 with anti-mouse Alexa Fluor 647, 4G8 with anti-mouse Alexa Fluor 488 and 11A5 with anti-mouse Alexa Fluor 488). 3D stacks were acquired following the same parameters for confocal laser scanning (magnification, resolution, laser power, offset and dwell time). All subfields were acquired from the same slide. All segmentations of the protein inclusions were done on Imaris 9.5.1 and 9.6 with the surface module following the same procedure for each marker (surface detail, absolute intensity, thresholds) (Fig, 1A). After segmentation, all volumes of the resulting objects (3D reconstructions) were summed up. The final score (%) represents the percentage of the volume of the stack covered by the selected marker staining.

**Figure 1.**
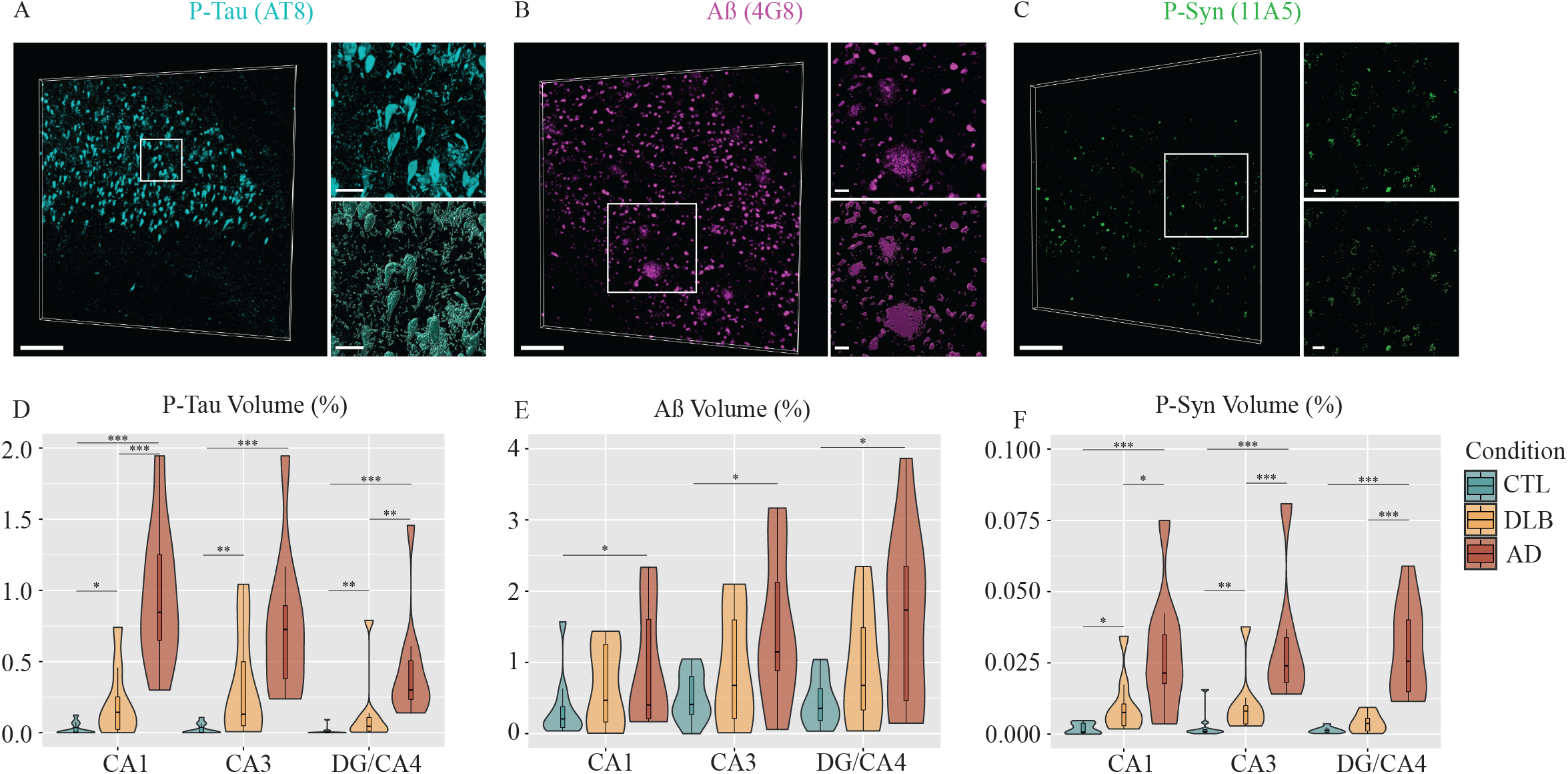
P-Tau, Aß and P-Syn loads show distinct disease-associated, subregional patterns in the hippocampus of AD and DLB patients. P-Tau, Aß, and P-Syn burdens were assessed via immunofluorescence staining (big panels and zoom in small upper panels) and volumetric analysis (volume percentage of 3D z-stack acquisition) in CA1, CA3 and DG/CA4 of age-matched CTLs, AD and DLB-mixed AD patients. The staining signal was segmented (with Imaris 9.6) (small lower panels) and their volume measured from 3D high-resolution acquisitions of each hippocampal subfield for each sample. (A) Paired helical filaments and neurofibrillary tangles of P-Tau (cyan) were stained with anti-AT8, example of an 80-year-old male AD patient (case 21). (B) Aß plaques and subcellular inclusions (magenta) were stained with anti-4G8, example of a 90-year-old male AD patient (case 30). (C) P-Syn inclusions, Lewy bodies and Lewy neurites (green) were stained with anti-11A5, example of a 64-year-old male DLB patient (case 15). Violin plots (with box plots) show that (D) P-Tau levels are significantly higher in the AD samples as compared to both DLB and CTLs in all three hippocampal subfields, but a significant increase was also observed in DLB samples compared to age-matched CTLs. (E) Aß levels are significantly higher in AD samples in all three subfields compared to CTLs, whereas no significant differences are seen between DLB and both AD and CTL. Aß loads follow the same subregional pattern in DLB but with lower values. (F) P-Syn loads are significantly increased in AD in all subfields, and in CA1 and CA3 in DLB compared to age-matched controls. P-Syn volumes are also significantly higher in AD samples compared to DLB. Wilcoxon-Mann-Whitney U-test p-values are indicated in the graphs: \**P* < 0.05; \*\**P* < 0.01 and \*\*\**P* < 0.001. Scale bars in A, B and C big panels = 200 µm, small panels = 30 µm.

#### MIC-MAC 2.0

Microglia and Immune Cells Morphologies Analyser and Classifier (MIC-MAC) 2.0 is a software that allows for automatic segmentation of microglia and immune cells from fluorescent microscopy 3D acquisition. With respect to the previous version^68^, automation and time performance have been improved.

#### New Automated Segmentation of Microglia

The image processing pipeline is now fully automated. While the previous version was based on a semi-supervised approach, meaning that a portion of the acquisition was needed to be labelled as positive signal or background, the new version automatically segments structures of interest by tuning parameters related to the image processing pipeline. A flowchart of the image processing pipeline is presented in Supplementary Fig. 2. The algorithm works by elaborating the 2D images of the 3D stacks one by one. First, a noise filtering is applied ^69^ with sigma = 25, followed by a background subtraction phase made with a gaussian filter (sigma = [10 - 50]). After an intensity adjustment [matlab::imadjust], the image is the deconvolved by using the Lucy-Richardson method^70,71^. Finally, positive pixels are detected via global thresholding by defining a parameter for the threshold t. The only two parameters that have been changed throughout this phase are sigma from the gaussian filter and t from the global thresholding. The parameters have been tuned accordingly to the amount of noise present in the image, which can change due to numerous factors (e.g. microscope parameterization, tissue conservation and antibody penetration). All morphologies of microglia were further validated by overlaying the segmented microglia object on the original 3D stack with the Iba1 staining.

#### Feature extraction and selection

Automated morphological feature extraction is performed at cell resolution^68^. Out of 62 morphological features and derived measures from the first version of MIC-MAC, we have selected only a subset of 16 features by discarding highly correlated features (Pearson, higher than 80%). Among the 16 kept features are morphological and graph-based measures, that all represent a certain aspect of the 3D morphology and help to measure fine changes and alterations. A detailed description for each of the 16 features can be found in Supplementary Fig. 3 with an illustration of prototypical minimum and maximum extreme values.

#### Artifact Removal

Segmentation artifacts were automatically removed in three different steps: (1) By thresholding cell volume and keeping cells whose volume is between ∼48,000 and ∼216,000 µm^3^, similar to values already present in literature. The volumetric range was visually validated. (2) By considering the amount of cell body surface that is included in the border of the acquisition. This allows to remove cells whose shape is incomplete and thus may alter the results. We specifically remove cells whose shared surface with the border is bigger than 7.5% of the total cell surface. This threshold has been manually tuned by visual validation. (3) By training a classifier based on morphological features. We trained a classifier (Boosted Tree – Ada Boost) ^72^ via manual labelling of artifacts and applied the classification to the whole dataset with k-fold validation to prevent overfitting. The removal via cell volume is performed right after the image processing segmentation in order to speed up the feature extraction phase. The latter two, are performed after the feature extraction phase right before the analysis.

#### Cluster analysis

All 16 selected features were initially normalized to the range 0 to 1 over the complete set of cells. We used Ward hierarchical clustering ^73^ on the set of 16 features. All the features have been normalized by range into a mapping from 0 to 1 renderings of cells belonging to specific clusters at different heights of the dendrogram. We assessed that *n*=7 was the best set-up to segregate distinct morphological clusters. These differences were further validated by the statistical analysis of features distribution among clusters. Cells were subsequently projected on a Uniform Manifold Approximation and Projection (UMAP)^74^ for visualization purpose.

### Statistical tests

For statistical comparisons between populations, we used Mann-Whitney U Test. For evaluating score and p-value of correlations we used Spearman Correlation. All tests were performed at a significance level of 5 %. In accordance with the explorative nature of the analysis, no correction for multiple testing was necessary.

### Data availability

Source code and raw data will be made publicly available after the peer-revision process.

## Results

### Co-occurrence and severity patterns of Tau, Aß and Synuclein pathologies in the hippocampal subfields of AD and DLB samples

To characterize the differences between Tau, Aß and Synuclein pathologies across AD, DLB and age-matched CTLs, we quantified the volumetric distribution of P-Tau, Aß and P-Syn using high-resolution confocal microscopy. The analysis of the fluorescent staining of AT8 for P-Tau and 4G8 for Aß antibodies, which are typically used for neuropathology assessment, both showed expected inclusions and labeled structures, i.e. typical paired helical filaments (PHF) and NFT with AT8^75^, Aß plaques and some amyloid precursor protein (APP) detected in neuronal soma with 4G8^76^ (Fig. 1A-B; Supplementary Fig.1B). The 11A5 P-Syn staining revealed different types of inclusions, from Lewy neurite-like structures to intracellular granular staining (Fig. 1C; Supplementary Fig.1B). To validate the 11A5 antibody against PS129 α-Synuclein, we immunostained 3 μm-thick serial FFPE sections of the amygdala from a Lewy body positive DLB case with 11A5, 81A (Millipore) and EP1536Y (Abcam) antibodies all directed against the same target PS129. All antibodies revealed a similar distribution pattern associated with Synuclein pathology such as Lewy neurites, Lewy bodies and intracellular vacuolar aggregations in neurons (Supplementary Fig. 4A). We then immunostained 100 μm-thick sections of PFA-fixed subiculum-entorhinal cortex of the same case, with a combination of an anti-PS129 antibodies (11A5, 81A or EP1536Y), an antibody against pan-axonal neurofilament to detect neurons (NF-H) and DRAQ7™ to stain nuclei and analyzed the 3D confocal images (Supplementary Fig. 4B). Again, all stainings between PS129 antibodies were broadly similar with labeled structures resembling Lewy bodies and Lewy neurites found enclosed in neurons double-stained for NF-H and DRAQ7™. The 11A5 positive structures found in our cohort of AD and DLB samples, also resembled Lewy neurites and Lewy bodies but also vacuolar aggregations and PHF-like, which were also seen with the 81A antibody (Supplementary Fig. 5).

After validation of the 11A5 antibody, we acquired representative 3D image stacks of AT8, 4G8, or 11A5 staining of the CA1, CA3 and DG/CA4 subfields for each case. The images were segmented with Imaris software 9.6 to estimate the relative volume covered by the pathology of interest (Fig. 1 A-C). We found variations in the pathological protein loads across samples and conditions that were generally compatible with the neuropathology staging of the samples. However, we observed all three P-Tau, Aß, and P-Syn pathologies in AD as well as in DLB cases. The majority of DLB cases were already described with partial AD pathologies. The age-matched CTLs showed no or low level of P-Tau staining and low to moderate Aß labelling in neuronal-like soma and in few extra-cellular deposits. In our CTLs group, we additionally observed very few P-Syn intra-cellular inclusions in cells looking like pyramidal neurons but no Lewy bodies or Lewy neurite like structures. The AD group typically showed the highest levels for all pathologies in the three hippocampal subfields. The DLB group showed low to intermediate levels of P-Tau, Aß, and P-Syn accumulations (Fig. 1 D-F).

More precisely, P-Tau burdens were significantly higher in all subfields in AD and DLB compared to the CTLs (AD vs CTLs: *P* < 0.001, DLB vs CTLs: *P <* 0.05) and followed a similar subregional pattern across conditions with the highest loads found in CA1 and CA3 and lowest in DG/CA4. The severity of the Tau pathology was considerably higher with the most extreme values in CA1 AD, and DG/CA4 AD compared to DLB respective subfields (*P* < 0.001 in CA1; *P* < 0.05 in DG/CA4). We found a distinct subregional pattern of Aß loads, again, conserved across the conditions and CTLs with the highest Aß burdens in DG/CA4 and CA3 and lowest in CA1. Only Aß AD values were significantly higher than CTLs ones for all subfields (*P <* 0.05). The subregional distribution of P-Syn loads appeared more uniform across subfields but with the highest loads found in CA1 and CA3 in AD and DLB. To be noted, the volume recovered by P-Syn staining were very low compared to Aß of P-Tau ones. We found statistical significances for all subfields compared to CTLs for AD (*P* < 0.001), and for CA1 (*P <* 0.05) and CA3 *(P <* 0.01) in DLB. Surprisingly, the P-Syn burdens were much higher in AD than DLB samples for all subfields (*P <* 0.05 CA1, *P <* 0.001 CA3 and DG/CA4). Our data suggest that Tau, Aß and Synuclein pathologies follow a specific subregional hippocampal pattern that is shared across AD and DLB. However, burdens were more severe in AD, with CA1 AD particularly affected by P-Tau accumulations.

Next, we examined the association between the three pathologies across subfields and conditions. Positive associations between P-Tau and Aß and between P-Syn and Aß volumes of respectively R^2^_Spear_ = 0.30 and 0.26 were found across subfields and conditions but differ subregionally with superior values in the CA1 (respectively R^2^_Spear_ = 0.42 and 0.37 *(P <* 0.05)) and a loss of statistical significance in the CA3 and DG/CA4. However, we found a strong positive correlation between P-Tau and P-Syn volumes across conditions (R^2^_Spear_ = 0.79; *P* < 0.001) that was preserved in all subfields (CA1: R^2^_Spear_ = 0.75; CA3: R^2^_Spear_ = 0.79; DG/CA4: R^2^_Spear_ = 0.78; *P* < 0.001) (Fig. 2A-B). To better define P-Tau and P-Syn associations at the cellular level, we tested double staining of P-Tau with a PSer422 rabbit antibody to pair with P-Syn (11A5, mouse) and DRAQ7™ (Fig. 2C). We confirmed that P-Tau PSer422 (rabbit) staining colocalized with AT8 ones in a CA1 AD case (Fig. 2D). We observed a global co-distribution of P-Tau and P-Syn in the same microenvironment, however, colocalizations in the same neuron were sparse and mono-stained neurons more common. Thus, the strong positive correlation between P-Tau and P-Syn does not point out intertwined intracellular co-pathologies but rather simultaneous mono P-Tau or P-Syn accumulations in neighboring hippocampal neurons.

**Figure 2.**
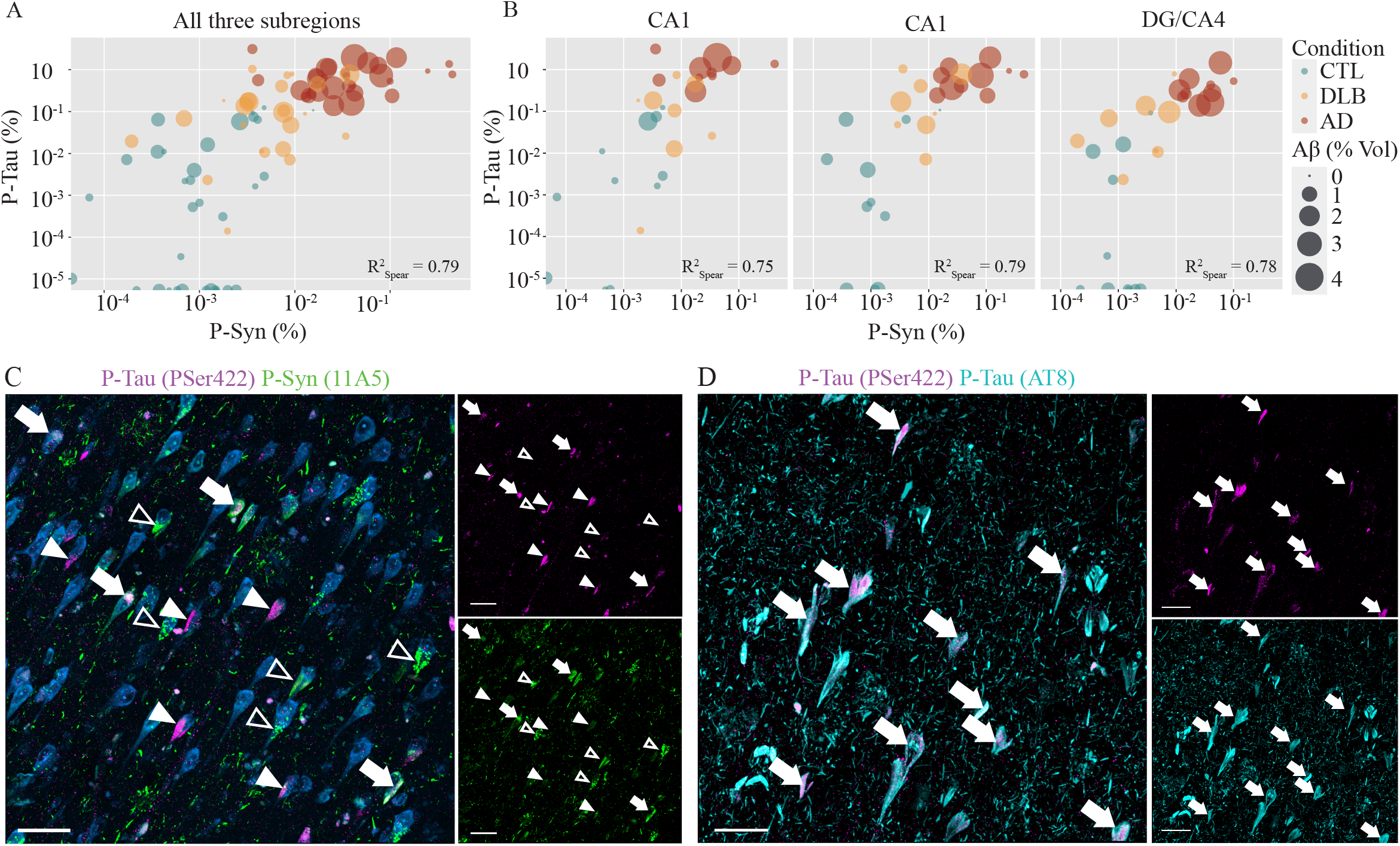
P-Tau and P-Synuclein loads are strongly correlated across hippocampal subregions and conditions. The Spearman correlation bubble charts (log_2_ scale) display a strong positive correlation between P-Tau and P-Syn loads across (A) conditions and subfields (R^2^_Spear_ = 0.79) and (B) at subregional level: CA1 (R^2^_Spear_ = 0.75), CA3 (R^2^_Spear_ = 0.79) and DG/CA4 (R^2^_Spear_ = 0.78). Corresponding Aß loads are labeled by the size of the bubble. Lower but positive correlation are observed between P-Tau and Aß (R^2^_Spear_ = 0.3) and between P-Syn and Aß (R^2^_Spear_ = 0.26) across the three subregions. (C) The correlation between P-Tau and P-Syn is also visible at cellular level with mainly mono-positive neurons for one or the other type of inclusion, but also some rare double-positive neurons that contain P-Tau and P-Syn inclusions at the same time. CA1 from an 82-year-old male AD patient (case 22) immunostained for P-Syn (mouse, 11A5, green), P-Tau (rabbit, PSer422, Thermofisher 44-764G, magenta) and fluorescent DNA dye DRAQ7™ (blue). Rare pyramidal neurons that co-express P-Syn and P-Tau are indicated by full arrowheads. More commonly, pyramidal neurons are positive for either P-Tau (full triangles) or P-Syn (empty triangles) inclusions. To facilitate visualization of mono-respectively double-positive neurons, the location of arrows and triangles is kept in all staining images. Nuclei of all cells are stained in blue. (D) Colocalization (full arrows) of AT8 (mouse, cyan) and PSer422 (rabbit, magenta) is confirmed in the CA1 of a 90-year-old AD patient (case 28). Scale bars (C, D): 30 μm.

### Microglia morphological responses in hippocampal subfields of AD and DLB patients

To assess the subregional microglia responses that paralleled the local pathological context, we took advantage of a pipeline for image analysis, previously published, that allows for semi-automated segmentation and classification of microglia cells based on their morphologies^68^. Indeed, morphologies of microglia partially reflect their responses; surveying microglia are usually very ramified, whereas activated microglia present an amoeboid-like morphology. For the purpose of this study that requires a time-consuming processing, we first implemented to our microglia morphological classifier called Microglia and Immune Cells Morphologies Analyser and Classifier (MIC-MAC)^68^ a 2.0 version which offers numerous improvements. MIC-MAC 2.0 allows a fully automated, more precise and faster segmentation of large 3D confocal stacks, an automated discard of the stack-border microglia that could create artifact structures and a simplified analysis relying on 16 morphological features that recapitulate in detail the core parameters of microglia 3D morphology (Supplementary Fig. 2 and 3). For this study, we captured more than 35,000 individual microglia across 87 3D stacks, representative of each of the three hippocampal subfields for each case of our cohort. Before extraction of the 3D features of microglia, we found a comparable density of Iba1 staining across subfields and conditions (Supplementary Fig. 6). After segmentation, MIC-MAC 2.0 imposed a volume threshold to select individual microglia but discard microglia accumulations that appeared as merged structures, such as the plaque associated microglia, from the analysis.

### Variations of microglia morphological features across hippocampal subfields in AD and DLB cases

First, we measured the 16 morphological features of extracted microglia (list and definition in Supplementary Fig. 3) to assess the precise changes in their morphologies across subfields and conditions. Interestingly, we found that DLB values present a trend similar to AD samples for almost all features and subfields, but without reaching significance. Among the 16 features, we found only four of them significantly different in AD compared to CTLs in at least one subfield. The compactness was the only feature significantly increased in all subfields in AD compared to CTL *(P <* 0.05 for CA1 and CA3, *P <* 0.001 for DG/CA4) (Fig. 3A). Volume/number of edges significantly increased, and node density decreased in CA1 (both *P* < 0.05) and DG AD (respectively *P* < 0.05 and *P <* 0.01) (Fig. 3B-C). Ending nodes density increased in CA3 AD *(P <* 0.05) (Fig. 3D). Intra-condition, we found that the feature average node degree is significantly higher in CA1 AD than in CA3 and DG/CA4 AD *(P <* 0.05 and *P <* 0.05; Fig. 3E), S-metric and node density lower in CA3 than DG in DLB (*P <* 0.05 Fig. 3F, and *P <* 0.05 Fig. 3C). Overall, compared to the CTLs microglia from the same subfield, CA1 AD microglia adopted a general morphology with higher compactness and loss of branching. AD DG/CA4 microglia followed a similar trend but to a lesser degree. AD CA3 microglia also showed a decrease in their complexity but with a distinct fine-tuning of their morphology.

**Figure 3.**
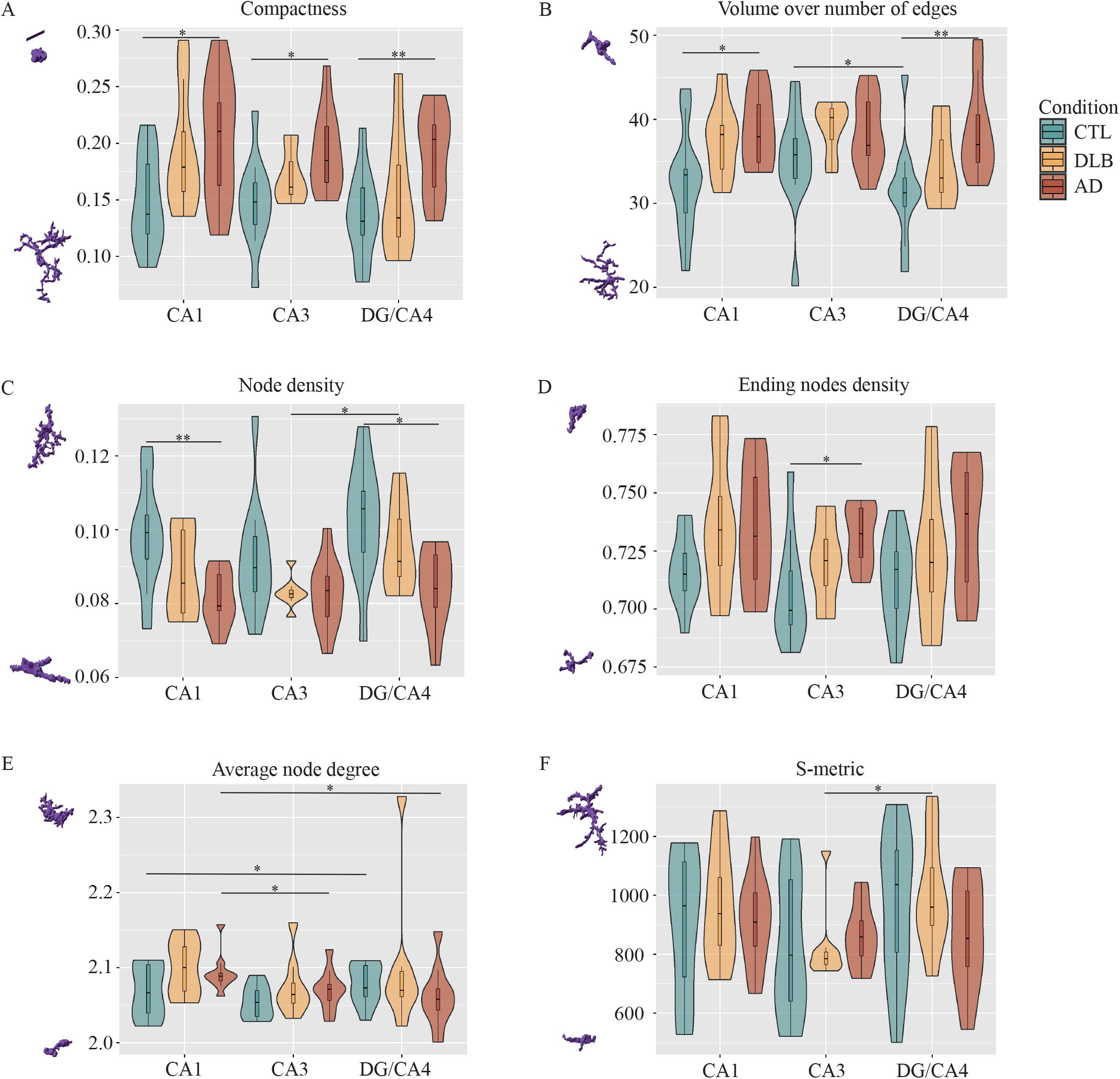
Subregional morphological alterations of microglia in the hippocampus of AD patients. Using MIC-MAC 2.0, we have extracted 16 geometrical features per cell of more than 35,000 individual microglia collected in CA1, CA3 and DG/CA4 subfields from AD, DLB and age-matched controls. None of the analyzed microglia morphological features was significantly changed in the DLB samples but follow a trend similar to the AD condition. All the following reported changes are observed in AD vs CTL samples. (A) Compactness is significantly increased in all three hippocampal subregions, (B) and volume over number of edges is significantly increased in CA1 and DG/CA4. (C) Node density significantly decreases in CA1 and DG/CA4, as does ending nodes density in CA3 (D). Two features, namely average node degree and s-metric present significant intra-condition regional changes. Wilcoxon-Mann-Whitney U-test p-values are indicated in the graphs: \**P* < 0.05; \*\**P* < 0.01 and \*\*\**P* < 0.001. Scaled prototypic morphologies with highest and lowest value for the selected feature have been added on the left side. Scale bar = 20µm.

### Microglia morphological clusters composition in the hippocampus AD and DLB patients

Features analysis allowed us to dissect the fine details of microglia morphological changes but does not reflect on how microglia populations are composed at a local level. To further characterize how microglia populations are remodeled across the hippocampal subfields in AD and DLB, we applied a cluster analysis to our matrix of morphological data. Briefly, we have selected seven clusters after a Ward hierarchical clustering based on a normalized set of the 16 features (Fig. 4A). The seven clusters show distinct morphological patterns and specific characteristics and were distributed accordingly on an UMAP with prototypical morphologies (Fig. 4B and Supplementary Fig. 7A). Briefly, clusters 1, 2 and 7 are composed by branched cells with high node density, volume and polarity. Cluster 3 represents the most compact and smallest microglia. Clusters 4, 5 and 6 show intermediate shapes mainly segregated by distinct features such as volume (higher in cluster 5), mean edge length (higher in cluster 4, not shown), and node density (higher in cluster 6) (Supplementary Fig. 7B-G). We then analyzed clusters relative abundances in CTLs, AD, and DLB. We found a general decrease of the most ramified clusters 1 and more severely of cluster 7 in AD compared to CTLs (respectively *P* < 0.05 and *P* < 0.001) and in DLB for 7 (*P* < 0.05). Cluster 3 and 5 were the two clusters enriched in both disease conditions (Fig. 5A) (cluster 3: AD *P* < 0.001, DLB *P* < 0.01) (cluster 5: AD *P* < 0.001, DLB *P* < 0.01). Clusters 2 and 6 showed similar ratios in all conditions and CTLs but cluster 4 was slightly but significantly increased in AD vs CTLs (*P* < 0.05) (Fig. 4C). At the subregional level, all conditions taken together, we found cluster 4 in lower quantity in DG/CA4 compared to CA3 (*P* <0.01) (Supplementary Fig. 8). When we analyzed the changes of cluster composition by subfield across conditions, we reached significance for cluster 3 increase for all subfields in AD compared to CTLs (*P* < 0.05 in CA1; *P* < 0.01 in CA3 and *P* < 0.05 in DG/CA4) and in CA3 DLB *(P* < 0.05); for cluster 5 increase in CA1 AD, DG/CA4 AD (*P* < 0.01) and CA3 DLB *(P* < 0.05), and for cluster 7 decrease in CA1 and DG/CA4 AD (*P* < 0.05) (Fig. 4D). Overall, we observe a tight regulation of the morphological cluster composition across subfields, with an enrichment of the two potential “disease” subgroups in AD, especially in the CA1 and DG/CA4, and a depletion of the ramified cluster 7. In DLB, alterations appeared more profound in the CA3 with the cluster 5 increase balancing cluster 7 decrease. The stability of cluster 2 and 6 across conditions imply that some subgroups of microglia do not react to the abnormal protein accumulations. The morphological cluster composition highlights the microglia functional heterogeneity. Some clusters specifically changed depending on the subfield and condition, and others were seemingly unreactive to the pathological context. This analysis further implies a subregional influence on microglia at the hippocampus level. In the next step, we investigated how these subfield specificities could tie up to the pathological context.

**Figure 4.**
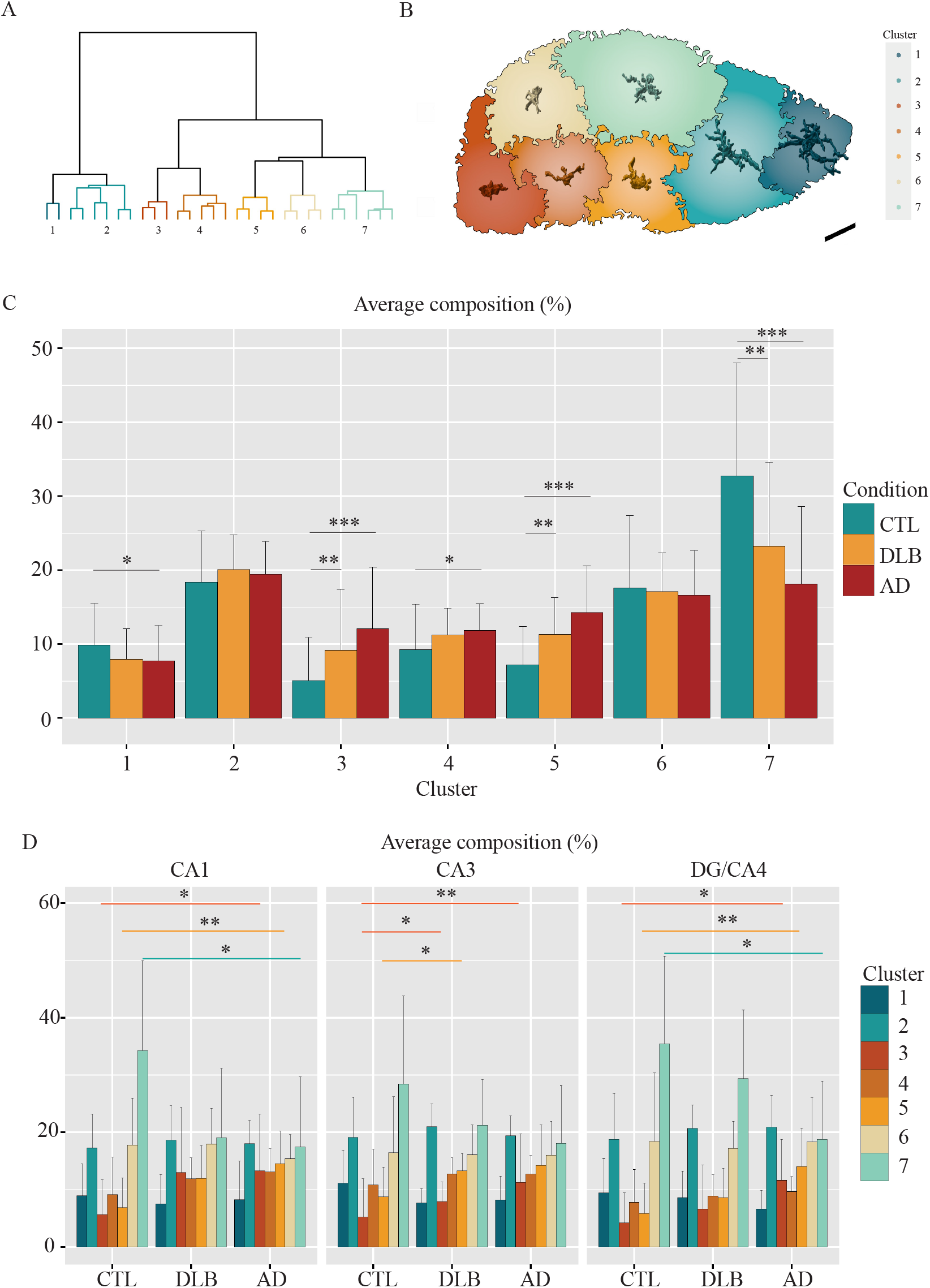
Microglia morphological cluster distribution varies according to the disease and the subfield. Ward hierarchical clustering on the matrix of 35,000 microglia based on 16 features defines seven distinct morphological clusters. (A) Hierarchical clustering dendrogram. (B) Scaled prototypical morphologies of microglia associated to the seven clusters inside the UMAP. (C-D) Histograms showing the variation in abundance (average composition in %) of the seven clusters according to the condition (C) and across subfield per condition (D). Overall, clusters 3 and 5 are enriched in the disease conditions, whereas cluster 7 is depleted. Scale bar in B = 30µm. Wilcoxon-Mann-Whitney U-test p-values are indicated in the graphs: \**P* < 0.05; \*\**P* < 0.01 and \*\*\**P* < 0.001.

**Figure 5.**
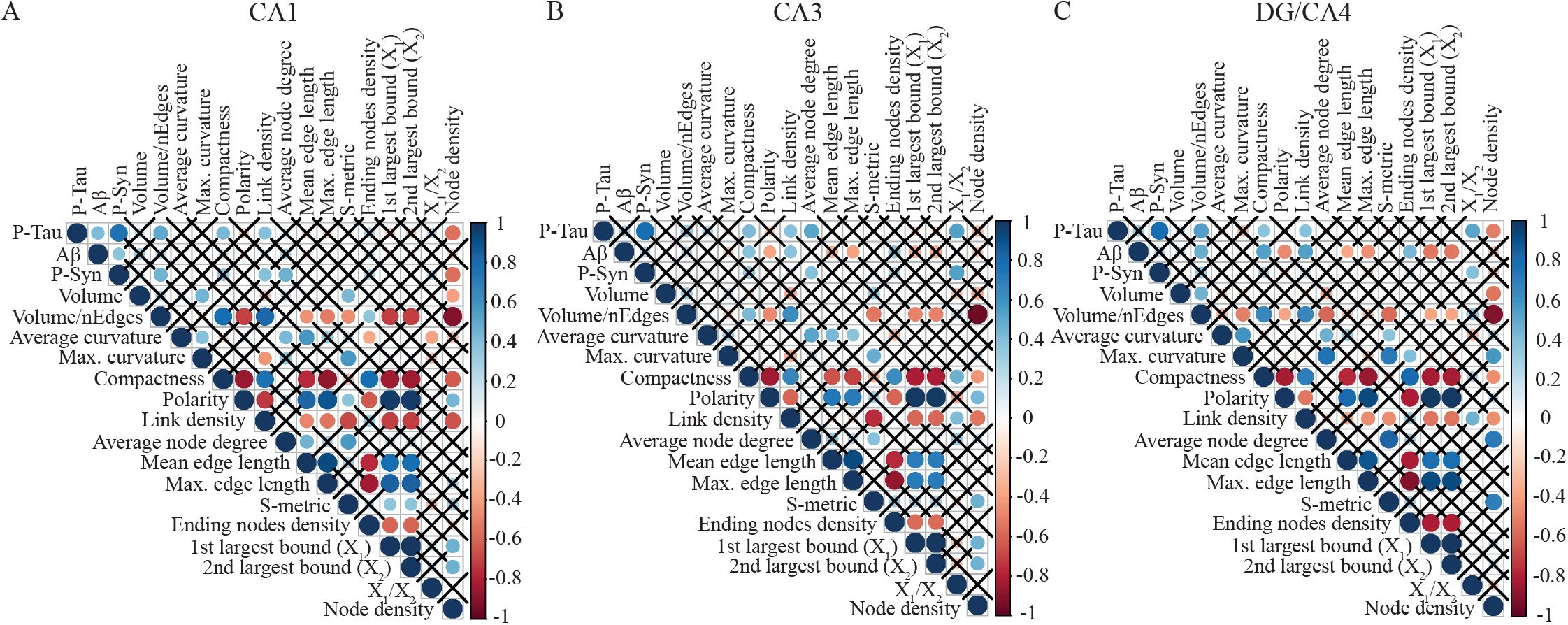
Microglial morphological features correlate with P-Tau, Aß, and P-Syn burdens in the hippocampus and follow a subregional pattern. Correlation plots between microglial morphological features and pathologies in CA1 (A) and CA3 (B), and DG/CA4 (C) in CTL, AD and DLB conditions. A list and description of all morphological features with prototypic morphologies for lowest and highest value can be found in S.Fig 5. Various microglia morphological changes are correlated to one or two proteinopathies in a subregion-dependent manner. Statistically significant values (Spearman’s correlation) are indicated by a dot, which color gives an estimation of the correlation (positive correlations in blue, negative correlations in red). Black crosses indicate unsignificant correlations.

### Associations between morphological microglia changes and P-Tau, Aß and P-Syn loads in AD and DLB hippocampi

To further estimate how each pathology could modulate microglia responses in the hippocampus, we first evaluated the associations (statistically significant) between morphological features and P-Tau, Aß and P-Syn burdens across subfields (Fig 5). Interestingly, the associations of P-Tau, Aß and P-Syn with features changed across subfields. In the CA1, we found significant correlations of some of the most altered features in AD with P-Tau (4) and/or P-Syn (4), most of the time in combinations at similar correlation values. In the CA3, some features were significantly associated with P-Tau (4) in pairs with Aß (2) or P-Syn (2), and 3 features uniquely to Aß. In DG/CA4, a higher range of features were positively or negatively associated with Aß (8), sometimes in pairs with P-Tau (3), with P-Tau alone (1) and P-Tau in pairs with P-Syn (1). The compactness, a feature statistically increased in AD, was associated with P-Tau loads in all subfields, but in pair with Aß only in the CA3 and DG/CA4. Volume/number of edges (Nedges) also changed in AD CA1 and DG/CA4 showed an association with both P-Tau and P-Syn in the CA1, and with P-Tau and Aß in DG/CA4. This data implies that P-Tau in association with P-Syn pathology triggered the most significant changes in the CA1, however Aß when highly expressed is a more dynamic modifier of microglia morphology.

At the cluster level, we also found a strong effect of the subregion on the associations between pathologies and clusters (Fig 6). First, we observed similar intercluster associations across the subfield. The increased “disease” cluster in AD and DLB, 3 and 5, are the most strongly positively associated in CA1 with this association preserved in CA3 and DG/CA4. Cluster 3 maintains a negative correlation with cluster 1 and 7 in all subfields. Cluster 5 stays positively correlated with cluster 4 in all subfields and negatively associated with cluster 7 in all subfields. Cluster 7, the largest subgroup in CTLs, seems to be depleted in favor of clusters 3 and 5, with cluster 3 potentially partially resulting of cluster 1 transformation. Cluster 5, poorly ramified, could also represent a transitional form toward cluster 3, more amoeboid-like, but stays in higher proportion across disease and subfields. Again, the stability of cluster 2 and 6 may indicate non-reactive microglia subgroups to their pathological micro-environment. Then, we found that cluster relationship with P-Tau, Aß and P-Syn burdens varied among subfields. In CA1, we found positive correlations between cluster 5, and negative ones for cluster 7, and the pair of P-Tau and P-Syn loads, and positive one between cluster 4 and P-Syn. In CA3, cluster 3 was positively correlated with the pair of P-Tau and Aß loads, clusters 4 and 5 with P-Tau loads. In DG/CA4, cluster 3 was associated with Aß loads only and cluster 4 with P-Tau ones only, cluster 5 and 7 with the pair of P-Tau and Aß loads.

**Figure 6.**
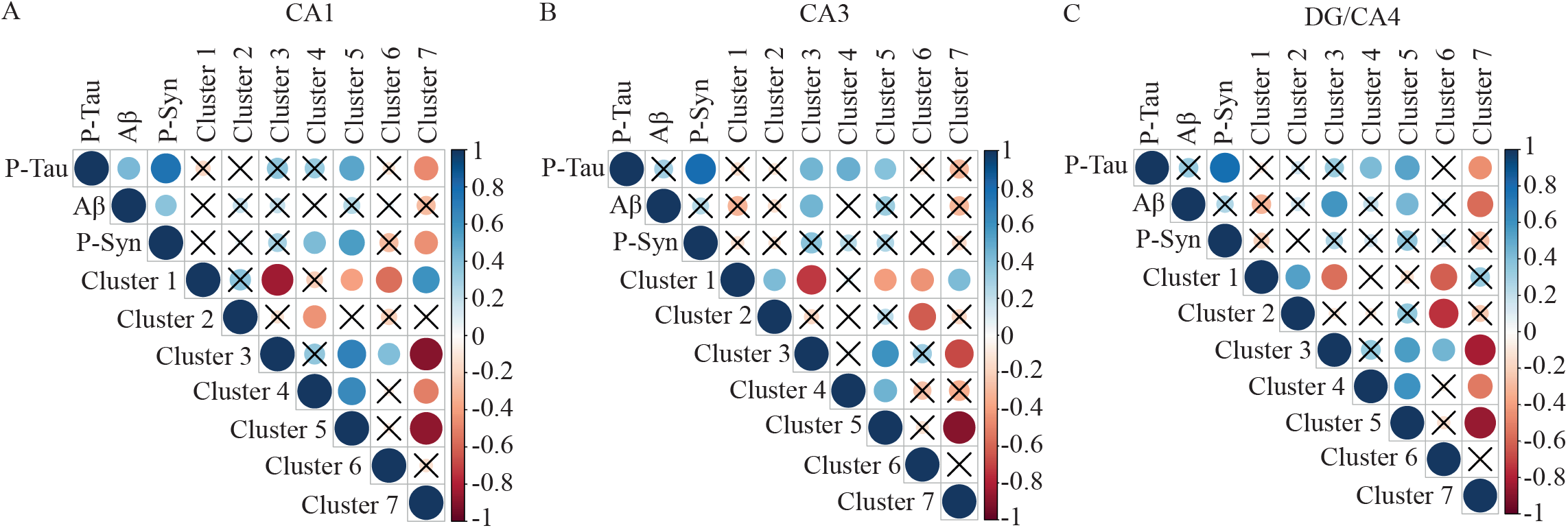
Microglial morphological clusters correlate with P-Tau, Aß, and P-Syn burdens in the hippocampus and follow a subregional pattern. Correlation plots between microglial morphological clusters and pathologies in CA1 (A) and CA3 (B), and DG/CA4 (C) of all conditions. In a subfield-dependent manner, disease-enriched clusters 3 and 5 are positively correlated with one or more pathologies whereas disease-depleted cluster 7 present negative correlations with proteinopathies. Statistically significant values (Spearman’s correlation) are indicated by a dot, which color gives an estimation of the correlation (positive correlations in blue, negative correlations in red). Black crosses indicate unsignificant correlations.

The strong associations between P-Tau, Aß and P-Syn pathologies and microglia morphologies suggest a tight inter-dependence of the pathological context and microglia responses. The severity, distribution but also co-occurrence of the three pathologies could impact microglia phenotypic heterogeneity and condition the level of their response.

## Discussion

Here we studied the subregional distribution and severity of the P-Tau, Aß and P-Syn pathologies along with the morphological changes of microglia in the hippocampal CA1, CA3 and DG/CA4 subfields across age-matched CTL, AD and DLB conditions. We have obtained high-content data about subregional patterning of pathologies in the human hippocampus across conditions and its relationship with the remodelling of the microglia local population.

### P-Tau, Aß and P-Syn burdens overlap and show specific subregional patterns in the hippocampus of AD and DLB cases

Here we report a high co-occurrence of the three types of pathologies studied, Tau, Aß, and Synuclein in hippocampal subfields of AD and DLB cases. P-Tau, Aß and P-Syn burdens were all more severe in AD. The concomitance of AD- and DLB-pathologies is commonly observed in age-related NDDs^77^ and reported in both AD and DLB ^24,27^ in numerous brain areas ^26,31,32,78^. However, our study revealed a complex tapestry of subregional pathological imprint in AD and DLB hippocampi, which each of the pathologies showing a specific subfield pattern. We found that P-Tau loads were more enriched toward CA1 and Aß ones toward DG/CA4. P-Syn loads were slightly higher in CA1/CA3 but appeared more homogeneous across subfields. DLB samples presented intermediate values of AD-typical and Synuclein pathologies. In this study, we have analyzed the medium part of the hippocampus, i.e., the body, between the uncus and *Corpus geniculatum laterale*, a region which is usually poorly described, contrary to the head at the uncus level or the tail close to the *Corpus geniculatum laterale*, both more regularly used for neuropathology diagnostic and research. We cannot exclude differences with previous neuropathological reports as it has been shown that the hippocampus is molecularly and functionally diverse along its longitudinal axis ^79,80^ and may be affected by different local level of pathologies. Finally, our quantification method, based on 3D confocal images, is more precise and reflects volumetric measures compared to the traditional surface measures. Our data are generally consistent with existing literature regarding the subregional distribution of Tau pathology in AD and DLB ^81,82^, but slightly differ regarding Aß subregional distribution ^83^ and P-Syn higher loads in AD. We also report low Aß and P-Syn accumulations but almost no P-Tau inclusions in our age-matched non-demented CTLs, which is in accordance with previous reports ^84–89^. Even though the P-Syn burdens that we have measured represent very low volumes compared to Aß and P-Tau ones, the use of high-resolution confocal microscopy allowed us to reveal frequent P-Syn intracellular granular inclusions in cells identified as pyramidal neurons. The higher P-Syn loads in AD cases compared to DLB seem at first counter-intuitive. We think that the hippocampus is relatively preserved of a more spread and severe Synuclein pathology as we have observed only a few Lewy body-like structures in our analysis. The typical accumulations of P-Syn into granules that we observed in cells identified as pyramidal neurons could represent an early or light stage of Synuclein pathology in the hippocampus. The few P-Syn granules found in CTLs samples could be in favor of such an hypothesis.

Interestingly, we reported stronger associations between P-Tau and P-Syn burdens than between P-Tau or P-Syn and Aß loads. By high-resolution confocal microscopy, we found that the positive correlation between Tau and Synuclein pathologies depended mainly on exclusive distribution of P-Tau or P-Syn inclusions in neighbouring hippocampal neuronal populations rather than on a colocalization in the same neuron that could suggest a systematic mechanistic association. However, we observed rare P-Tau and P-Syn double-positive hippocampal neurons carrying the two types of pathologies such as previously found in the subiculum ^90^, the amygdala ^91^ and parahippocampal gyri ^92^ in AD and DLB cases. In a previous study, the local co-distribution of NFTs and Lewy bodies was also found more frequent in limbic areas than other brain parts ^26^. How P-Syn- and P-Tau-bearing neighbouring neurons co-inhabit the AD and DLB hippocampus and how the two pathologies impact each other remains to be understood.

### Hippocampal microglia morphological remodelling is impacted by complex subregional pathological context

Morphology of microglia only partially reflect their responses. However, visual description is commonly used to obtain a rapid readout of their alterations in CNS diseases. Indeed, the homeostatic microglia harbor a highly ramified profile in contrast to activated microglia that appear smaller, amoeboid-like with no or low branches. Here, we have analyzed 3D morphologies to characterize the local reorganization of the microglia population in the hippocampus subregions of AD and DLB patients. We previously developed an AI pipeline called MIC-MAC, to classify and quantify morphologies of individual microglia labelled with the Iba1 marker in large 3D confocal image stacks of human or mouse postmortem brains ^68^. With MIC-MAC 2.0 presented here, we have significantly improved the automation and precision of the segmentation and added new features to discard artefacts upstream from the analysis. A caveat of our approach is that MIC-MAC 2.0 automatically discards all microglia accumulations, where the cells are so closely intermingled that our pipeline cannot separate them into individual cells after the segmentation process. In our hands, even super-resolution microscopy could not resolve the human microglia cellular boundaries in these accumulations. This argument strengthened our decision to focus the MIC-MAC analysis on individual cells to overcome potential contaminations in our data. Our analysis created a substantial amount of information with a sampling of 35,000 extracted microglia. By quantifying their 16 morphological features, our analysis highlighted conditions and subregional particularities. When compared to their respective subfield in CTLs, we found statistical significances only for the AD condition. However, DLB variation trends always mimic AD ones showing that the presence of concomitant proteinopathies already initiated local remodeling even if subtler. Many reports found low to mild microglial changes in DLB samples. These conclusions were supported by 2D morphology analyses and quantifications of some activation markers in the hippocampus^61^ and other brain areas such temporal lobe, the superior frontal gyrus^59^ and cerebral cortex^60^. However, our analysis implies that DLB microglia are not so different from AD ones in their ability to respond and that their alterations are associated with the level of severity of DLB and AD pathologies in the parenchyma. High compactness was the only predictive feature for AD across subfields, strengthening the observation that microglia tend to become more amoeboid in AD, even when not attached to the plaques. At the morphological clusters level, as in previous reports ^61,93^, we show a decrease of the most branched and tortuous subgroups and an increase of amoeboid-like clusters significantly in AD. Interestingly the disease-increased clusters are also found at low levels in CTLs and could be products of the ageing process or the onset of pathological mechanisms^94^. Our data also further suggest functional heterogeneity among microglia local subgroups. Indeed, we found that two morphological clusters stayed relatively stable over the conditions, while others were specifically depleted toward simplified morphologies. We have also uncovered two stereotypical morphologies enriched in AD and DLB, one amoeboid-like and one semi-ramified. It could reflect differences in response to pathological stressors and further illustrates the residual heterogeneity within the microglia population in health and disease^55^. Our study demonstrates that Tau, Aß and Synuclein pathologies impact the local microglia remodelling in the AD and DLB hippocampus. We found fine-tuned associations with specific features or clusters that reflect the tight interdependence of microglia states with its pathological environment. Microglia responses are often characterized in monogenic models where only one pathology is expressed, with these signatures often interpolated to the human condition. Many of the microglia AD signatures, such as the disease-associated microglia^95–97^, have been characterized in monogenic animal models mimicking Aß pathology and were often observed in microglia surrounding Aß plaques. If these signatures are fully or partially, regionally or sub-regionally recapitulated in human conditions still needs to be further characterized ^98–103^. Recently, Gerrits et al.^104^ have defined by single-nuclei RNA sequencing that some signatures of microglia were selectively associated with Aß, or with combined Aß and Tau pathologies, in the occipital and occipitotemporal cortices of AD patients. Such molecular heterogeneity of microglia is expected in the AD hippocampus. Indeed, we observed that hippocampal microglia alterations are most often associated with dual pathologies. Microglia local responses are tightly correlated with the co-occurrence and severity of Tau, Aß and Synuclein pathologies. It does not contradict our observation of the strong impact of Aß on microglia morphologies in the CA3, which has also been described in amyloidosis mouse models. Nevertheless, it emphasizes the dual role of Tau and Synuclein pathologies on exacerbating microglia responses, particularly in the AD CA1. The complex pathological pattern of the human condition certainly represents a factor of divergence of microglia phenotypes between AD patients and amyloidosis rodent models.

## Conclusion

Our analysis suggests that the co-occurrence of the three Tau, Aß and Synuclein pathologies is frequent in the hippocampus of AD and DLB patients with a general severity largely increased in AD cases. Each pathology seems to follow a distinct subregional pattern, mainly conserved across AD and DLB. Our study also implies a synergistic effect of the three pathologies on microglia transformations that tightly regulate the morphological composition of the local microglia population. The combination of high burdens of P-Tau and P-Syn with increased microglia alterations could participate in accelerated atrophy of the CA1 in AD.

## Abbreviations

Aß: Amyloid-ß
AD: Alzheimer’s Disease
APP: Amyloid precursor protein
CA: *Cornu ammonis*
CTL: Control
DG: *Gyrus dentatus*
DLB: Dementia with Lewy Bodies
MIC-MAC: Microglia and Immune Cells Morphologies Analyser and Classifier
PMD: Post-mortem delay
P-Syn: Hyperphosphorylated α-synuclein
P-Tau: Hyperphosphorylated tau
UMAP: Uniform manifold approximation and projection

## Acknowledgements

The authors thank the Douglas-Bell Brain Bank and its staff for providing human brain samples, the Bioimaging Facility of the Luxembourg Centre for Systems Biomedicine (LCSB) for support of microscopy, and the Reproducible Research Results (R3) team of the LCSB for promoting reproducible research. The 11A5 antibody was kindly provided by Dr. Manuel Buttini and Dr. Wagner Zago (Prothena Biosciences).

## Supplementary material

Supplementary material is available online.

## Authors contributions

DS.B conceived and supervised the study. S.F produced IHC on fixed tissue, S.F and DS.B performed confocal microscopy acquisitions and volumetric analysis. C.A designed MIC-MAC 2.0, with inputs from S.F, L.S and DS.B for validations. C.A and G.H produced statistical data analysis. O.U.H, C.S and JJ.G produced some support experiments. N.M provided human fixed samples. M.M provided FFPE samples and resources. A.S provided bioinformatic resources. S.F, C.A and G.H produced figures. S.F, C.A, M.M and DS.B wrote the manuscript. All authors contributed to the final version of the manuscript.

## Figure legends

**Supplementary Figure 1. Hippocampal subfields and confocal stainings of P-Tau, Aß and P-Syn across conditions and hippocampal subfields**. (A) Anatomically defined CA1, CA3 and DG/CA4 subfields of the hippocampus for proteinopathy and microglia 3D z-stack acquisitions. Hippocampal section of an 83-year-old female AD patient (case 24) with Aß (4G8, green) and microglia (Iba1, magenta) stainings. (B) 50-100µm thick hippocampal sections were immunostained with the AT8 antibody against P-Tau (Ser202, Thr205) (cyan), the 4G8 antibody against Aß (AA17-24) (magenta) and the 11A5 antibody against P-Syn (Ser129) (green). The stainings show heterogenous distribution and types of inclusions across conditions and hippocampal subregions. Scale bars in (A) = 1000 µm, in (B) = 100 µm.

**Supplementary Figure 2 MIC-MAC 2.0 analysis workflow**. MIC-MAC 2.0 pipeline with illustrations of acquisition (A), automated image processing (B), manual validation via overlay (C), feature extraction (D), feature analysis (E) and cluster analysis (F). (A-C) Original Iba1 staining of microglia is represented in white, segmented microglia cells in magenta. (D) Microglia 3D geometrical features are based on nodes (connection points) and edges (branches). (F) Spatial cluster color coded overlay. Scale bars in (A-C), (F) = 100µm.

**Supplementary Figure 3. Morphological features with minimum and maximum values**. List and description of all morphological features with prototypic morphologies for lowest and highest feature value. Significant changes based on Wilcoxon-Mann-Whitney U-test observed in AD samples are indicated with one (*P* < 0.05) or two arrows (*P* < 0.01) on the right side. Scale bars = 20µm.

**Supplementary Figure 4. Validation of P-Syn stainings with three different antibodies on FFPE samples of a neuropathologically confirmed DLB case**. 3-µm thick paraffin block sections from the amygdala and 50-100 µm thick sections from fixed hippocampus were obtained from the same 91-year-old male DLB patient (case 20) and were stained against P-Syn using three different antibodies, namely 11A5, 81A and ab51253, that all three recognize the P-Ser-129 epitope. (A) Sections from paraffin blocks were stained against P-Syn (11A4, 81A and ab51253, brown) based on the DAB/HRP substrate system. The lower rows represent a zoom of the indicated region in the upper row. (B) Sections from fixed samples were stained by immunofluorescence against P-Syn (11A4, 81A and ab51253; green), neurofilaments (NF-H, magenta) and all nuclei (DRAQ7™, blue). Scale bars in (A) upper row = 500 µm and lower row = 50 µm; (B) upper row = 100 µm and lower row = 10 µm.

**Supplementary Figure 5. Confocal description of P-Syn staining in a neuropathologically confirmed DLB case**. 50-100 µm thick sections from fixed hippocampus from a 90-year-old male AD patient (case 28) were immunostained against P-Syn (11A5, 81A; green), neurofilaments (NF-H, magenta) and all nuclei (DRAQ7, blue). The upper row shows the hippocampus (*stratum oriens* at left bottom corner) at 20x magnification, and the lower row represents a zoom of the marked areas in the pyramidal layer. P-Syn inclusions a present under various forms, including PHF-like (full triangle), Lewy neurite (empty triangle) and vacuolar aggregations (arrowhead). Scale bars upper row = 50 µm and lower row = 20 µm.

**Supplementary Figure 6. Iba1 density in hippocampal subfields of AD and DLB do not differ from age-matched CTLs**. Iba1 volume (%) across conditions and hippocampal subfields.

**Supplementary Figure 7. Microglia morphologies can be divided in seven clusters**. (A) Projection on a UMAP of the seven clusters where each of the more than 35,000 individual microglia is represented by a dot. (B-D) Distribution spectrum for volume, compactness and polarity on UMAP. (E-G) Violin plots showing the enrichment on volume, compactness and polarity in specific clusters.

**Supplementary Figure 8. Cluster distribution varies among hippocampal subregions**. Histogram showing the variation in abundance (average composition in %) of the seven clusters according to CA1, CA3 and DG/CA4 subregions. Only cluster 4 presents sa significant change: a decrease in DG/CA4 compared to CA3. Wilcoxon-Mann-Whitney U-test p-values are indicated in the graphs: \**P* < 0.05; \*\**P* < 0.01 and \*\*\**P* < 0.001.

## References

1. Dubois B. The Emergence of a New Conceptual Framework for Alzheimer’s Disease. Journal of Alzheimer’s Disease. 2018;62(3). doi:10.3233/JAD-170536

2. Burgess N, Maguire EA, O’Keefe J. The human hippocampus and spatial and episodic memory. Neuron. 2002;35(4):625–641. doi:10.1016/S0896-6273(02)00830-9

3. Bird CM, Burgess N. The hippocampus and memory: Insights from spatial processing. Nature Reviews Neuroscience. 2008;9(3):182–194. doi:10.1038/nrn2335

4. Lisman J, Buzsáki G, Eichenbaum H, Nadel L, Rangananth C, Redish AD. Viewpoints: How the hippocampus contributes to memory, navigation and cognition. Nature Neuroscience. 2017;20(11):1434–1447. doi:10.1038/nn.4661

5. Foo H, Thalamuthu A, Jiang J, et al. Associations between Alzheimer’s disease polygenic risk scores and hippocampal subfield volumes in 17,161 UK Biobank participants. Neurobiology of Aging. 2021;98:108–115. doi:10.1016/j.neurobiolaging.2020.11.002

6. Mrdjen D, Fox EJ, Bukhari SA, Montine KS, Bendall SC, Montine TJ. The basis of cellular and regional vulnerability in Alzheimer’s disease. Acta Neuropathologica. 2019;138(5):729–749. doi:10.1007/s00401-019-02054-4

7. Crist AM, Hinkle KM, Wang X, et al. Transcriptomic analysis to identify genes associated with selective hippocampal vulnerability in Alzheimer’s disease. Nature Communications. 2021;12(1). doi:10.1038/s41467-021-22399-3

8. Golomb J, Leon MJ, Kluger A, Tarshish C, Ferris SH, George AE. Hippocampal atrophy in normal aging: An association with recent memory impairment. Archives of Neurology. 1993;50(9). doi:10.1001/archneur.1993.00540090066012

9. Ferrarini L, Van Lew B, Reiber JHC, et al. Hippocampal atrophy in people with memory deficits: Results from the population-based IPREA study. International Psychogeriatrics. 2014;26(7). doi:10.1017/S1041610213002627

10. Basu J, Siegelbaum SA. The corticohippocampal circuit, synaptic plasticity, and memory. Cold Spring Harbor Perspectives in Biology. 2015;7(11). doi:10.1101/cshperspect.a021733

11. Yassa MA, Stark CEL. Pattern separation in the hippocampus. Trends in Neurosciences. 2011;34(10). doi:10.1016/j.tins.2011.06.006

12. Coras R, Pauli E, Li J, et al. Differential influence of hippocampal subfields to memory formation: Insights from patients with temporal lobe epilepsy. Brain. 2014;137(7). doi:10.1093/brain/awu100

13. Seok JW, Cheong C. Functional dissociation of hippocampal subregions corresponding to memory types and stages. Journal of Physiological Anthropology. 2020;39(1). doi:10.1186/s40101-020-00225-x

14. Sabattoli F, Boccardi M, Galluzzi S, Treves A, Thompson PM, Frisoni GB. Hippocampal shape differences in dementia with Lewy bodies. NeuroImage. 2008;41(3). doi:10.1016/j.neuroimage.2008.02.060

15. Kerchner GA, Deutsch GK, Zeineh M, Dougherty RF, Saranathan M, Rutt BK. Hippocampal CA1 apical neuropil atrophy and memory performance in Alzheimer’s disease. NeuroImage. 2012;63(1). doi:10.1016/j.neuroimage.2012.06.048

16. La Joie R, Perrotin A, De La Sayette V, et al. Hippocampal subfield volumetry in mild cognitive impairment, Alzheimer’s disease and semantic dementia. NeuroImage: Clinical. 2013;3. doi:10.1016/j.nicl.2013.08.007

17. Apostolova LG, Mosconi L, Thompson PM, et al. Subregional hippocampal atrophy predicts Alzheimer’s dementia in the cognitively normal. Neurobiology of Aging. 2010;31(7). doi:10.1016/j.neurobiolaging.2008.08.008

18. Adler DH, Wisse LEM, Ittyerah R, et al. Characterizing the human hippocampus in aging and Alzheimer’s disease using a computational atlas derived from ex vivo MRI and histology. Proceedings of the National Academy of Sciences of the United States of America. 2018;115(16). doi:10.1073/pnas.1801093115

19. Zhao W, Wang X, Yin C, He M, Li S, Han Y. Trajectories of the hippocampal subfields atrophy in the alzheimer’s disease: A structural imaging study. Frontiers in Neuroinformatics. 2019;13. doi:10.3389/fninf.2019.00013

20. Lashuel HA. Rethinking protein aggregation and drug discovery in neurodegenerative diseases: Why we need to embrace complexity? Current Opinion in Chemical Biology. 2021;64:67–75. doi:10.1016/J.CBPA.2021.05.006

21. Braak H, Braak E. Neuropathological stageing of Alzheimer-related changes. Acta Neuropathologica. 1991;82(4). doi:10.1007/BF00308809

22. Thal DR, Rüb U, Orantes M, Braak H. Phases of Aβ-deposition in the human brain and its relevance for the development of AD. Neurology. 2002;58(12):1791–1800. doi:10.1212/WNL.58.12.1791

23. McKeith IG, Galasko D, Kosaka K, et al. Consensus guidelines for the clinical and pathologic diagnosis of dementia with Lewy bodies (DLB): Report of the consortium on DLB international workshop. Neurology. 1996;47(5):1113–1124. doi:10.1212/WNL.47.5.1113

24. McKeith IG, Boeve BF, Dickson DW et al. McKeith IG, Boeve BF, Dickson DW, et al. Diagnosis and management of dementia with Lewy bodies: Fourth consensus report of the DLB Consortium. Neurology. 2017;89(1).

25. Golde TE, Borchelt DR, Giasson BI, Lewis J. Thinking laterally about neurodegenerative proteinopathies. Journal of Clinical Investigation. Published online 2013. doi:10.1172/JCI66029

26. Colom-Cadena M, Gelpi E, Charif S, et al. Confluence of α-synuclein, tau, and β-amyloid pathologies in dementia with Lewy bodies. Journal of Neuropathology and Experimental Neurology. Published online 2013. doi:10.1097/NEN.0000000000000018

27. Robinson JL, Lee EB, Xie SX, et al. Neurodegenerative disease concomitant proteinopathies are prevalent, age-related and APOE4-associated. Brain. Published online 2018. doi:10.1093/brain/awy146

28. Irwin DJ, Grossman M, Weintraub D, et al. Neuropathological and genetic correlates of survival and dementia onset in synucleinopathies: a retrospective analysis. The Lancet Neurology. 2017;16(1). doi:10.1016/S1474-4422(16)30291-5

29. Mak E, Donaghy PC, McKiernan E, et al. Beta amyloid deposition maps onto hippocampal and subiculum atrophy in dementia with Lewy bodies. Neurobiology of Aging. 2019;73. doi:10.1016/j.neurobiolaging.2018.09.004

30. Twohig D, Nielsen HM. α-synuclein in the pathophysiology of Alzheimer’s disease. Molecular Neurodegeneration. 2019;14(1). doi:10.1186/s13024-019-0320-x

31. Ferreira D, Przybelski SA, Lesnick TG, et al. β-Amyloid and tau biomarkers and clinical phenotype in dementia with Lewy bodies. Neurology. 2020;95(24). doi:10.1212/WNL.0000000000010943

32. Schumacher J, Gunter JL, Przybelski SA, et al. Dementia with Lewy bodies: association of Alzheimer pathology with functional connectivity networks. Brain. Published online 2021. doi:10.1093/brain/awab218

33. Márquez-Ropero M, Benito E, Plaza-Zabala A, Sierra A. Microglial Corpse Clearance: Lessons From Macrophages. Frontiers in Immunology. 2020;11. doi:10.3389/fimmu.2020.00506

34. Lee CYD, Landreth GE. The role of microglia in amyloid clearance from the AD brain. Journal of Neural Transmission. 2010;117(8). doi:10.1007/s00702-010-0433-4

35. Luo W, Liu W, Hu X, Hanna M, Caravaca A, Paul SM. Microglial internalization and degradation of pathological tau is enhanced by an anti-tau monoclonal antibody. Scientific Reports. 2015;5. doi:10.1038/srep11161

36. Sanchez-Mejias E, Navarro V, Jimenez S, et al. Soluble phospho-tau from Alzheimer’s disease hippocampus drives microglial degeneration. Acta Neuropathologica. 2016;132(6). doi:10.1007/s00401-016-1630-5

37. Paolicelli RC, Jawaid A, Henstridge CM, et al. TDP-43 Depletion in Microglia Promotes Amyloid Clearance but Also Induces Synapse Loss. Neuron. 2017;95(2). doi:10.1016/j.neuron.2017.05.037

38. Španić E, Langer Horvat L, Hof PR, Šimić G. Role of Microglial Cells in Alzheimer’s Disease Tau Propagation. Frontiers in Aging Neuroscience. 2019;11. doi:10.3389/fnagi.2019.00271

39. Zeisel A, Moz-Manchado AB, Codeluppi S, et al. Cell types in the mouse cortex and hippocampus revealed by single-cell RNA-seq. Science. 2015;347(6226). doi:10.1126/science.aaa1934

40. Grubman A, Choo XY, Chew G, et al. Transcriptional signature in microglia associated with Aβ plaque phagocytosis. Nature Communications. 2021;12(1). doi:10.1038/s41467-021-23111-1

41. Bouvier DS, Murai KK. Synergistic actions of microglia and astrocytes in the progression of Alzheimer’s disease. Journal of Alzheimer’s Disease. 2015;45(4):1001–1014. doi:10.3233/JAD-143156

42. Sarlus H, Heneka MT. Microglia in Alzheimer’s disease. Journal of Clinical Investigation. 2017;127(9):3240–3249. doi:10.1172/JCI90606

43. De Strooper B, Karran E. The Cellular Phase of Alzheimer’s Disease. Cell. 2016;164(4). doi:10.1016/j.cell.2015.12.056

44. Walker LC. Aβ Plaques. Free neuropathology. 2020;1. doi:10.17879/freeneuropathology-2020-3025

45. Huang Y, Happonen KE, Burrola PG, et al. Microglia use TAM receptors to detect and engulf amyloid β plaques. Nature Immunology. 2021;22(5). doi:10.1038/s41590-021-00913-5

46. Keren-Shaul H, Spinrad A, Weiner A, et al. A Unique Microglia Type Associated with Restricting Development of Alzheimer’s Disease. Cell. 2017;169(7):1276-1290.e17. doi:10.1016/j.cell.2017.05.018

47. Krasemann S, Madore C, Cialic R, et al. The TREM2-APOE Pathway Drives the Transcriptional Phenotype of Dysfunctional Microglia in Neurodegenerative Diseases. Immunity. 2017;47(3). doi:10.1016/j.immuni.2017.08.008

48. Maphis N, Xu G, Kokiko-Cochran ON, et al. Reactive microglia drive tau pathology and contribute to the spreading of pathological tau in the brain. Brain. 2015;138(6). doi:10.1093/brain/awv081

49. Romero-Molina C, Navarro V, Sanchez-Varo R, et al. Distinct microglial responses in two transgenic murine models of TAU pathology. Frontiers in Cellular Neuroscience. 2018;12. doi:10.3389/fncel.2018.00421

50. Vautheny A, Duwat C, Aurégan G, et al. THY-Tau22 mouse model accumulates more tauopathy at late stage of the disease in response to microglia deactivation through TREM2 deficiency. Neurobiology of Disease. 2021;155. doi:10.1016/j.nbd.2021.105398

51. Choi I, Zhang Y, Seegobin SP, et al. Microglia clear neuron-released α-synuclein via selective autophagy and prevent neurodegeneration. Nature Communications. 2020;11(1). doi:10.1038/s41467-020-15119-w

52. George S, Rey NL, Tyson T, et al. Microglia affect α-synuclein cell-to-cell transfer in a mouse model of Parkinson’s disease. Molecular Neurodegeneration. 2019;14(1). doi:10.1186/s13024-019-0335-3

53. McQuade A, Blurton-Jones M. Microglia in Alzheimer’s Disease: Exploring How Genetics and Phenotype Influence Risk. Journal of Molecular Biology. 2019;431(9). doi:10.1016/j.jmb.2019.01.045

54. Salter MW, Stevens B. Microglia emerge as central players in brain disease. Nature Medicine. 2017;23(9):1018–1027. doi:10.1038/nm.4397

55. Masuda T, Sankowski R, Staszewski O, Prinz M. Microglia Heterogeneity in the Single-Cell Era. Cell Reports. 2020;30(5):1271–1281. doi:10.1016/j.celrep.2020.01.010

56. Prinz M, Jung S, Priller J. Microglia Biology: One Century of Evolving Concepts. Cell. 2019;179(2):292–311. doi:10.1016/j.cell.2019.08.053

57. Uriarte Huarte O, Richart L, Mittelbronn M, Michelucci A. Microglia in Health and Disease: The Strength to Be Diverse and Reactive. Frontiers in Cellular Neuroscience. 2021;15. doi:10.3389/fncel.2021.660523

58. Mackenzie IRA. Activated microglia in dementia with Lewy bodies. Neurology. 2000;55(1):132–134. doi:10.1212/WNL.55.1.132

59. Streit WJ, Xue QS. Microglia in dementia with Lewy bodies. Brain, Behavior, and Immunity. 2016;55:191–201. doi:10.1016/j.bbi.2015.10.012

60. Amin J, Holmes C, Dorey RB, et al. Neuroinflammation in dementia with Lewy bodies: a human post-mortem study. Translational Psychiatry. 2020;10(1). doi:10.1038/s41398-020-00954-8

61. Bachstetter AD, Van Eldik LJ, Schmitt FA, et al. Disease-related microglia heterogeneity in the hippocampus of Alzheimer’s disease, dementia with Lewy bodies, and hippocampal sclerosis of aging. Acta neuropathologica communications. 2015;3:32. doi:10.1186/s40478-015-0209-z

62. Surendranathan A, Su L, Mak E, et al. Early microglial activation and peripheral inflammation in dementia with Lewy bodies. Brain. 2018;141(12). doi:10.1093/brain/awy265

63. Chia R, Sabir MS, Bandres-Ciga S, et al. Genome sequencing analysis identifies new loci associated with Lewy body dementia and provides insights into its genetic architecture. Nature Genetics. 2021;53(3):294–303. doi:10.1038/s41588-021-00785-3

64. Montine TJ, Phelps CH, Beach TG, et al. National institute on aging-Alzheimer’s association guidelines for the neuropathologic assessment of Alzheimer’s disease: A practical approach. Acta Neuropathologica. 2012;123(1). doi:10.1007/s00401-011-0910-3

65. McKeith IG, Galasko D, Kosaka K, et al. Consensus guidelines for the clinical and pathologic diagnosis of dementia with Lewy bodies (DLB): Report of the consortium on DLB international workshop. Neurology. 1996;47(5). doi:10.1212/WNL.47.5.1113

66. Bouvier DS, Jones E V., Quesseveur G, et al. High Resolution Dissection of Reactive Glial Nets in Alzheimer’s Disease. Scientific Reports. Published online 2016. doi:10.1038/srep24544

67. Quesseveur G, Fouquier d’Hérouël A, Murai KK, Bouvier DS. A Specialized Method to Resolve Fine 3D Features of Astrocytes in Nonhuman Primate (Marmoset, Callithrix jacchus) and Human Fixed Brain Samples. In: Di Benedetto B, ed. Astrocytes: Methods and Protocols. Springer New York; 2019:85–95. doi:10.1007/978-1-4939-9068-9_6

68. Salamanca L, Mechawar N, Murai KK, Balling R, Bouvier DS, Skupin A. MIC-MAC: An automated pipeline for high-throughput characterization and classification of three-dimensional microglia morphologies in mouse and human postmortem brain samples. GLIA. 2019;67(8). doi:10.1002/glia.23623

69. Dabov K, Foi A, Katkovnik V, Egiazarian K. Image restoration by sparse 3D transform-domain collaborative filtering. In: Image Processing: Algorithms and Systems VI. Vol 6812.; 2008. doi:10.1117/12.766355

70. Richardson WH. Bayesian-Based Iterative Method of Image Restoration*. Journal of the Optical Society of America. 1972;62(1). doi:10.1364/josa.62.000055

71. Lucy LB. An iterative technique for the rectification of observed distributions. The Astronomical Journal. 1974;79. doi:10.1086/111605

72. Freund Y, Schapire RE. A Decision-Theoretic Generalization of On-Line Learning and an Application to Boosting. Journal of Computer and System Sciences. 1997;55(1). doi:10.1006/jcss.1997.1504

73. Ward JH. Hierarchical Grouping to Optimize an Objective Function. Journal of the American Statistical Association. 1963;58(301). doi:10.1080/01621459.1963.10500845

74. McInnes L, Healy J, Saul N, Großberger L. UMAP: Uniform Manifold Approximation and Projection. Journal of Open Source Software. 2018;3(29). doi:10.21105/joss.00861

75. Goedert M, Jakes R, Vanmechelen E. Monoclonal antibody AT8 recognises tau protein phosphorylated at both serine 202 and threonine 205. Neuroscience Letters. 1995;189(3):167–170. doi:10.1016/0304-3940(95)11484-E

76. Hunter S, Brayne C. Do anti-amyloid beta protein antibody cross reactivities confound Alzheimer disease research? Journal of Negative Results in BioMedicine. 2017;16(1):1–8. doi:10.1186/s12952-017-0066-3

77. Golde TE, Borchelt DR, Giasson BI, Lewis J. Thinking laterally about neurodegenerative proteinopathies. Journal of Clinical Investigation. Published online 2013. doi:10.1172/JCI66029

78. Walker L, McAleese KE, Thomas AJ, et al. Neuropathologically mixed Alzheimer’s and Lewy body disease: burden of pathological protein aggregates differs between clinical phenotypes. Acta Neuropathologica. 2015;129(5). doi:10.1007/s00401-015-1406-3

79. Strange BA, Witter MP, Lein ES, Moser EI. Functional organization of the hippocampal longitudinal axis. Nature Reviews Neuroscience. 2014;15(10). doi:10.1038/nrn3785

80. Ayhan F, Kulkarni A, Berto S, et al. Resolving cellular and molecular diversity along the hippocampal anterior-to-posterior axis in humans. Neuron. 2021;109(13):2091-2105.e6. doi:10.1016/j.neuron.2021.05.003

81. Braak H, Alafuzoff I, Arzberger T, Kretzschmar H, Tredici K. Staging of Alzheimer disease-associated neurofibrillary pathology using paraffin sections and immunocytochemistry. Acta Neuropathologica. 2006;112(4):389–404. doi:10.1007/s00401-006-0127-z

82. Coughlin DG, Ittyerah R, Peterson C, et al. Hippocampal subfield pathologic burden in Lewy body diseases vs. Alzheimer’s disease. Neuropathology and Applied Neurobiology. Published online 2020. doi:10.1111/nan.12659

83. D.R. T, U. R, M. O, H. B. Phases of Aβ-deposition in the human brain and its relevance for the development of AD. Neurology. 2002;58(12).

84. Crystal H, Dickson D, Fuld P, et al. Clinico-pathologic studies in dementia: Nondemented subjects with pathologically confirmed alzheimer’s disease. Neurology. 1988;38(11). doi:10.1212/wnl.38.11.1682

85. Markesbery WR, Jicha GA, Liu H, Schmitt FA. Lewy body pathology in normal elderly subjects. Journal of Neuropathology and Experimental Neurology. Published online 2009. doi:10.1097/NEN.0b013e3181ac10a7

86. Rodrigue KM, Kennedy KM, Devous MD, et al. β-amyloid burden in healthy aging: Regional distribution and cognitive consequences. Neurology. 2012;78(6). doi:10.1212/WNL.0b013e318245d295

87. Kovacs GG, Milenkovic I, Wöhrer A, et al. Non-Alzheimer neurodegenerative pathologies and their combinations are more frequent than commonly believed in the elderly brain: A community-based autopsy series. Acta Neuropathologica. Published online 2013. doi:10.1007/s00401-013-1157-y

88. Kawas CH, Kim RC, Sonnen JA, Bullain SS, Trieu T, Corrada MM. Multiple pathologies are common and related to dementia in the oldest-old: The 90 + Study. Neurology. Published online 2015. doi:10.1212/WNL.0000000000001831

89. Jansen WJ, Ossenkoppele R, Knol DL, et al. Prevalence of cerebral amyloid pathology in persons without dementia: A meta-analysis. JAMA - Journal of the American Medical Association. 2015;313(19). doi:10.1001/jama.2015.4668

90. Iseki E, Takayama N, Marui W, Uéda K, Kosaka K. Relationship in the formation process between neurofibrillary tangles and Lewy bodies in the hippocampus of dementia with Lewy bodies brains. Journal of the Neurological Sciences. 2002;195(1). doi:10.1016/S0022-510X(01)00689-X

91. Schmidt ML, Martin JA, Lee VMY, Trojanowski JQ. Convergence of Lewy bodies and neurofibrillary tangles in amygdala neurons of Alzheimer’s disease and Lewy body disorders. Acta Neuropathologica. 1996;91(5). doi:10.1007/s004010050454

92. Griffin WST, Liu L, Li Y, Mrak RE, Barger SW. Interleukin-1 mediates Alzheimer and Lewy body pathologies. Journal of Neuroinflammation. 2006;3:1–9. doi:10.1186/1742-2094-3-5

93. Paasila PJ, Davies DS, Kril JJ, Goldsbury C, Sutherland GT. The relationship between the morphological subtypes of microglia and Alzheimer’s disease neuropathology. Brain Pathology. 2019;29(6). doi:10.1111/bpa.12717

94. Candlish M, Hefendehl JK. Microglia Phenotypes Converge in Aging and Neurodegenerative Disease. Frontiers in Neurology. 2021;12. doi:10.3389/fneur.2021.660720

95. Keren-Shaul H, Spinrad A, Weiner A, et al. A Unique Microglia Type Associated with Restricting Development of Alzheimer’s Disease. Cell. 2017;169(7):1276-1290.e17. doi:10.1016/j.cell.2017.05.018

96. Krasemann S, Madore C, Cialic R, et al. The TREM2-APOE Pathway Drives the Transcriptional Phenotype of Dysfunctional Microglia in Neurodegenerative Diseases. Immunity. 2017;47(3). doi:10.1016/j.immuni.2017.08.008

97. Sala Frigerio C, Wolfs L, Fattorelli N, et al. The Major Risk Factors for Alzheimer’s Disease: Age, Sex, and Genes Modulate the Microglia Response to Aβ Plaques. Cell Reports. 2019;27(4). doi:10.1016/j.celrep.2019.03.099

98. Patir A, Shih B, McColl BW, Freeman TC. A core transcriptional signature of human microglia: Derivation and utility in describing region-dependent alterations associated with Alzheimer’s disease. Glia. 2019;67(7):1240–1253. doi:10.1002/glia.23572

99. Del-Aguila JL, Li Z, Dube U, et al. A single-nuclei RNA sequencing study of Mendelian and sporadic AD in the human brain. Alzheimer’s Research and Therapy. 2019;11(1). doi:10.1186/s13195-019-0524-x

100. Mathys H, Davila-Velderrain J, Peng Z, et al. Single-cell transcriptomic analysis of Alzheimer’s disease. Nature. 2019;570(7761). doi:10.1038/s41586-019-1195-2

101. Srinivasan K, Friedman BA, Etxeberria A, et al. Alzheimer’s Patient Microglia Exhibit Enhanced Aging and Unique Transcriptional Activation. Cell Reports. 2020;31(13). doi:10.1016/j.celrep.2020.107843

102. Chen WT, Lu A, Craessaerts K, et al. Spatial Transcriptomics and In Situ Sequencing to Study Alzheimer’s Disease. Cell. 2020;182(4):976-991.e19. doi:10.1016/j.cell.2020.06.038

103. Walker DG. Defining activation states of microglia in human brain tissue: an unresolved issue for Alzheimer’s disease. Neuroimmunology and Neuroinflammation. 2020;2020. doi:10.20517/2347-8659.2020.09

104. Gerrits E, Brouwer N, Kooistra SM, et al. Distinct amyloid-β and tau-associated microglia profiles in Alzheimer’s disease. Acta Neuropathologica. 2021;141(5):681–696. doi:10.1007/s00401-021-02263-w

