## Supplementary figures for "Concomitant AD and DLB pathologies shape subfield microglia responses in the hippocampus"

A

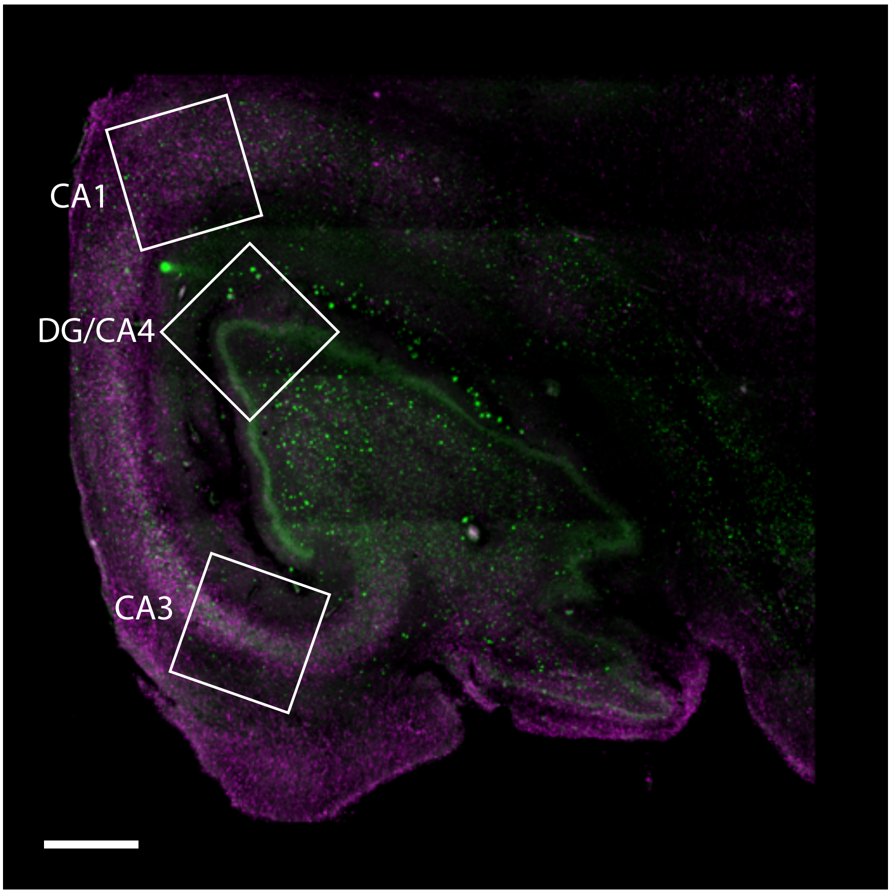

CA1
DG/CA4
CA3


B

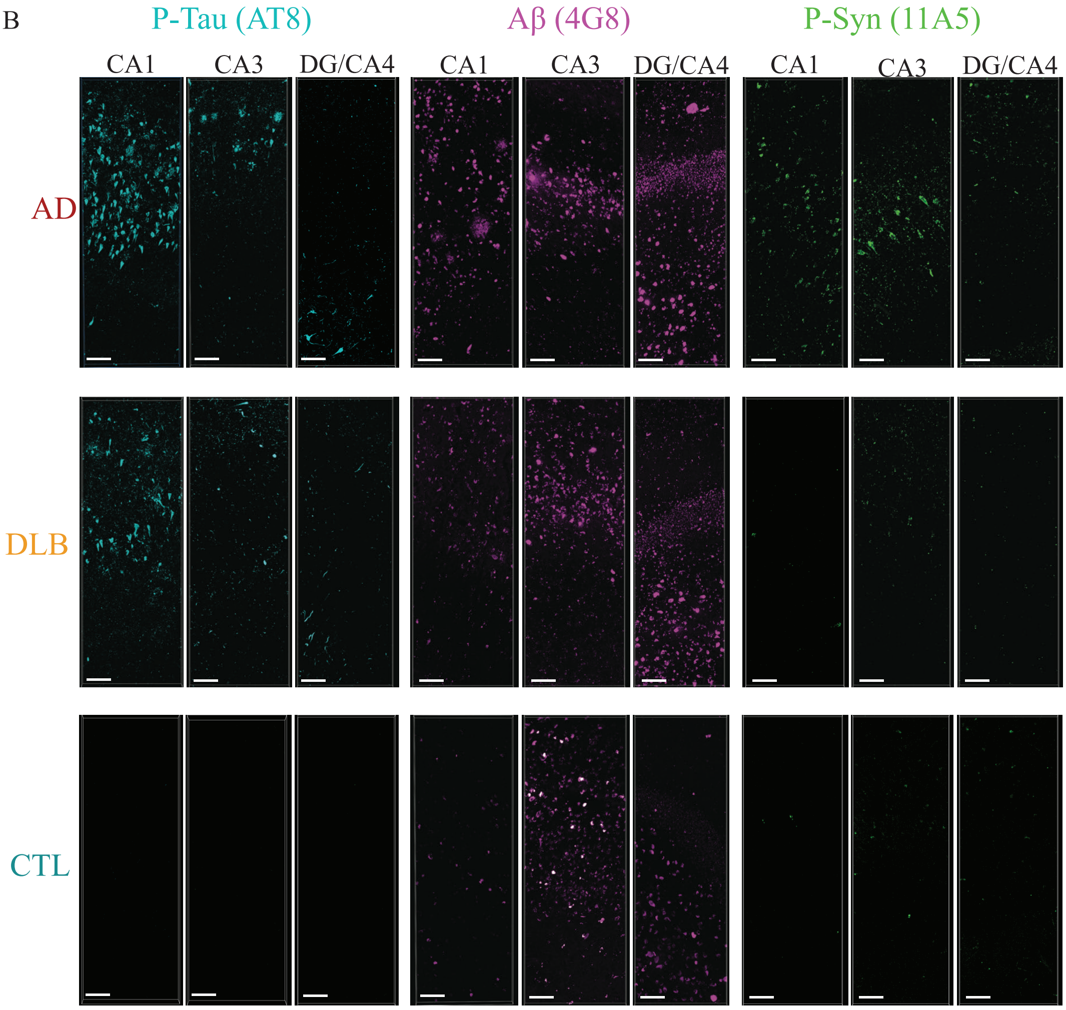

P-Tau (AT8)
Aβ (4G8)
P-Syn (11A5)
CA1
CA3
DG/CA4
CA1
CA3
DG/CA4
CA1
CA3
DG/CA4
AD
DLB
CTL


**SupFig.1**

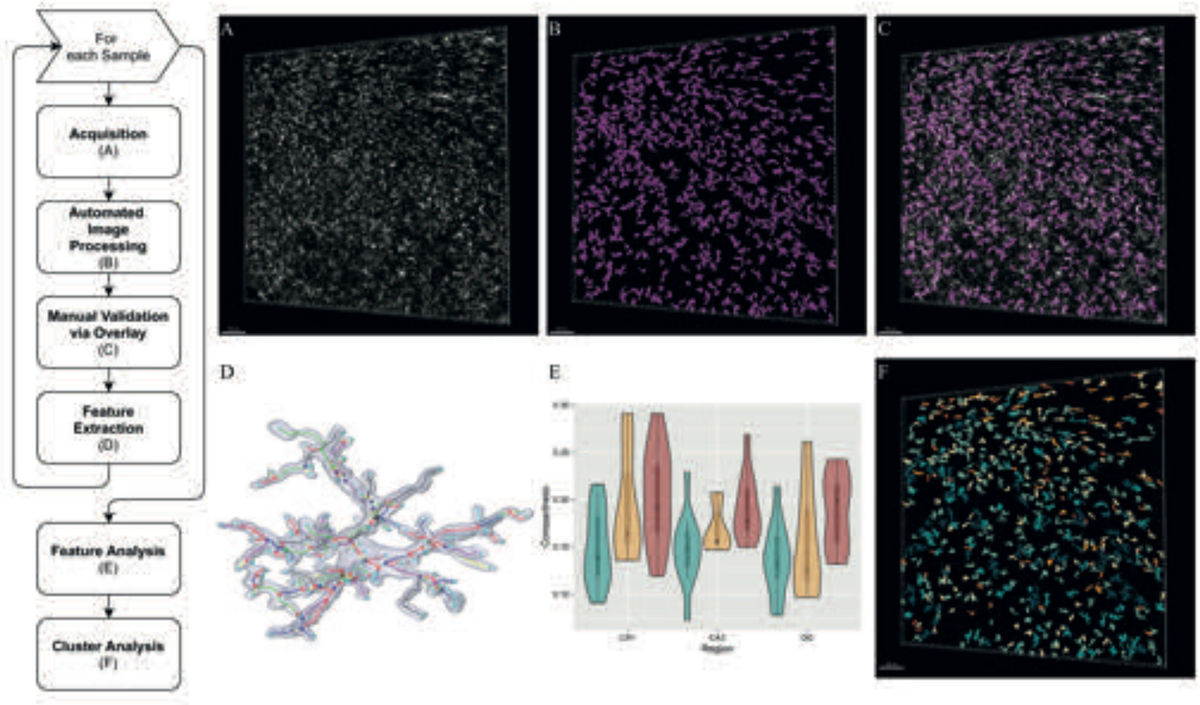
For each Sample
Acquisition (A)
Automated Image Processing (B)
Manual Validation via Overlay (C)
Feature Extraction (D)
Feature Analysis (E)
Cluster Analysis (F)
A
B
C
D
E
F

**SupFig 2**

| Feature<br>Description | Min value | Max value |
|---|---|---|
| **1st largest bound**<br>Largest dimension of bounding box enclosing a structure | | |
| **2nd largest bound**<br>2nd largest dimension of bounding box enclosing a structure | | |
| **1 over 2**<br>Ratio of 1st and 2nd lenghts (elongation) | | |
| **Average curvature**<br>Average curvature of projections (length of edge/distance of nodes) | | |
| **Average Node degree**<br>Average degree for all the nodes in the graph | | |
| **Compactness**<br>Ratio between structure volume and volume of convex hull | | |
| **Ending Node density**<br>Number of ending nodes over total number of nodes | | |
| **Link density**<br>Number of edges divided by the number of node pairs | | |
| **Max curvature**<br>Maximal curvature of projections (length of edge/distance of nodes) | | |
| **Max Edge Length**<br>Longest edge in the graph | | |
| **Mean Edge Length**<br>Mean length of all edges in the graph | | |
| **Node density**<br>Number of nodes divided by total volume | | |
| **Polarity**<br>Average direction of the projections emerging from the central node | | |
| **S-metric**<br>Summed products of nodal degrees across all edges | | |
| **Volume**<br>Number of voxels that constitute the volume | | |
| **Volume / number of Edges**<br>Volume in voxels divided by the number of existing edges | | |

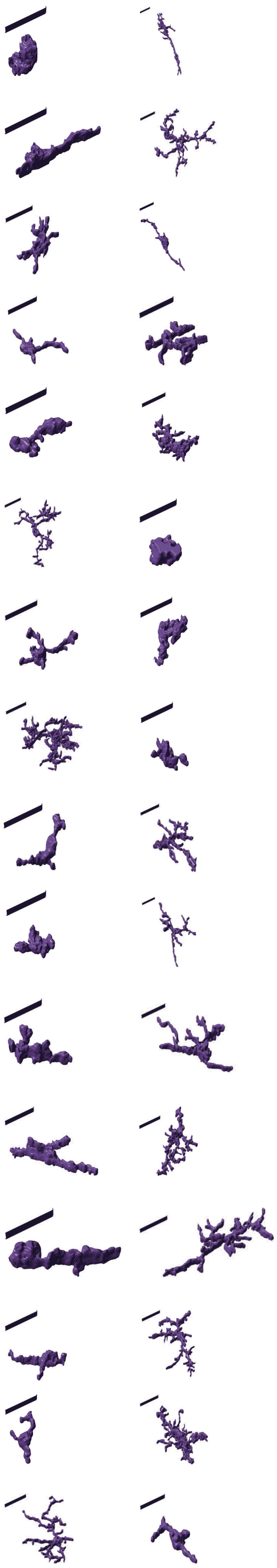

**SupFig 3**

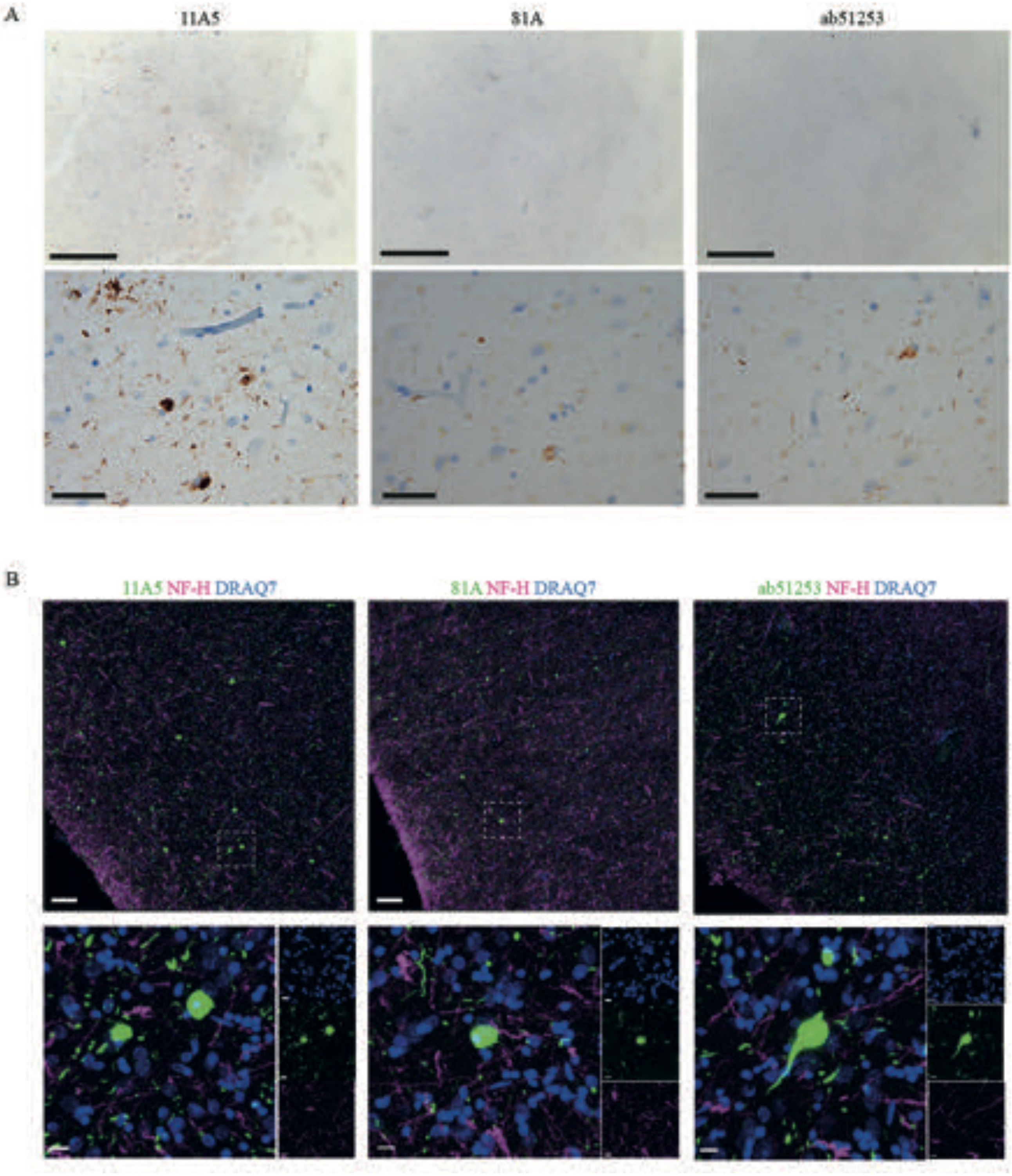
A
11A5
81A
ab51253
B
11A5 NF-H DRAQ7
81A NF-H DRAQ7
ab51253 NF-H DRAQ7

**SupFig 4**

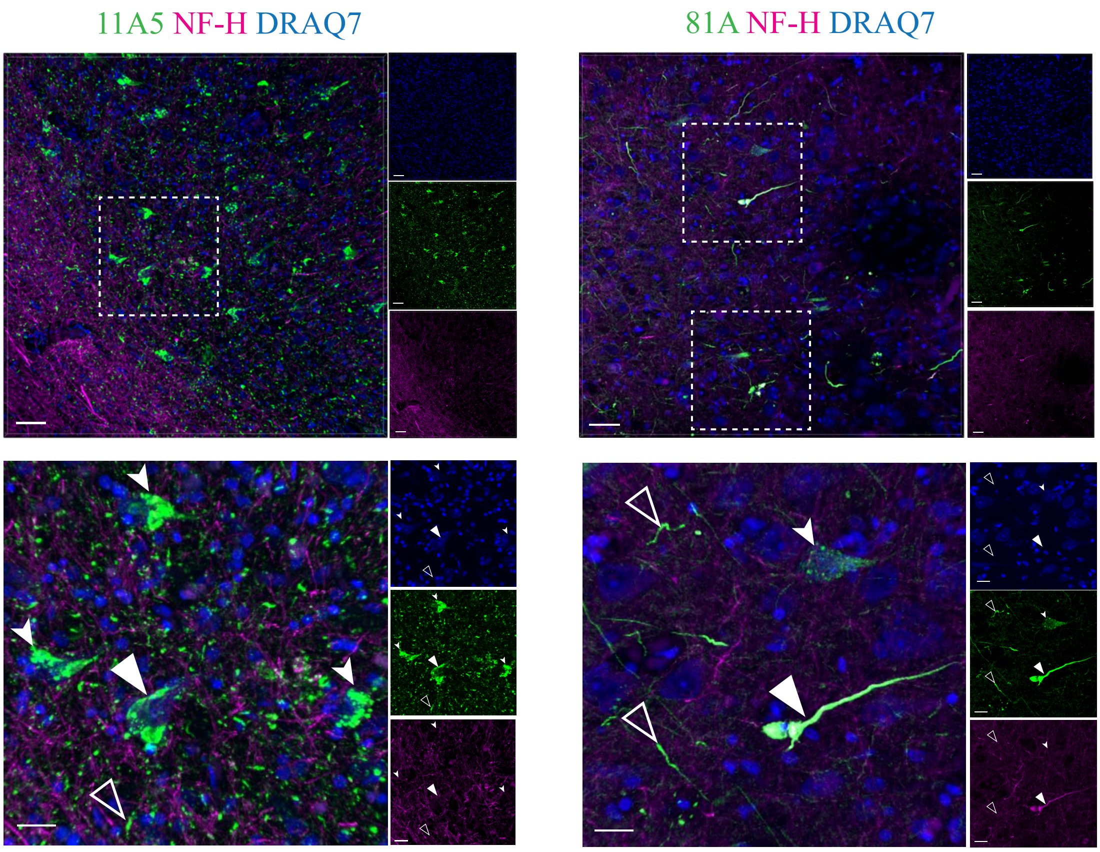


**SupFig 5**

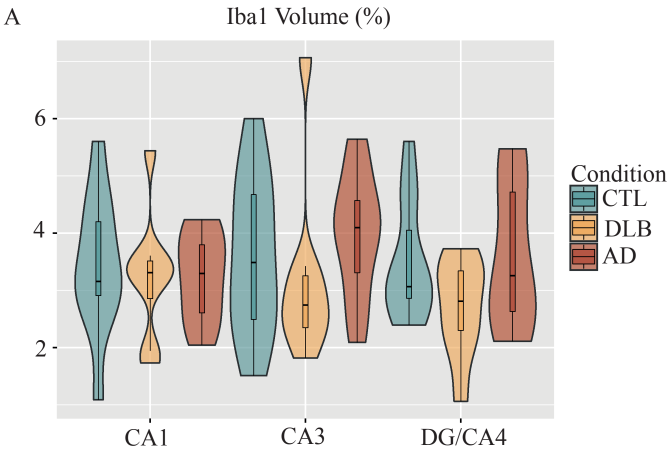


**SupFig 6**

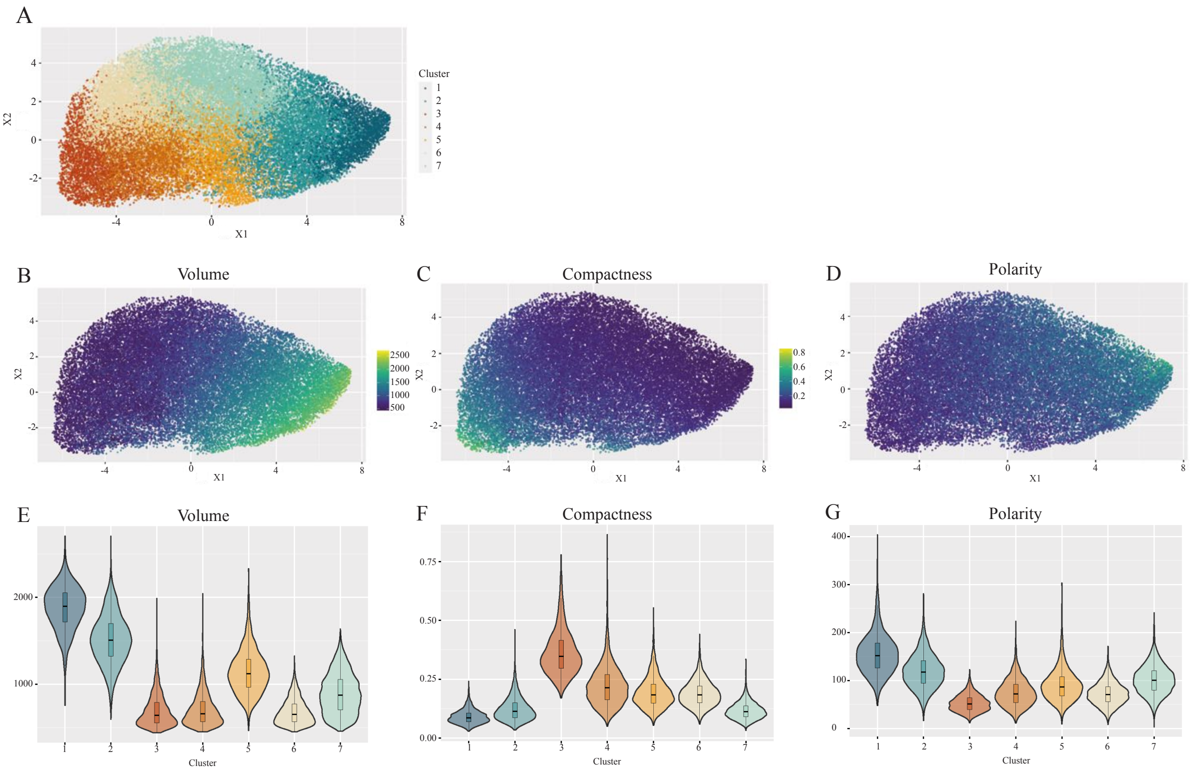
A
Cluster
1
2
3
4
5
6
7
X1
X2
B
Volume
C
Compactness
D
Polarity
E
Volume
F
Compactness
G
Polarity
Cluster

### Supp Fig 7

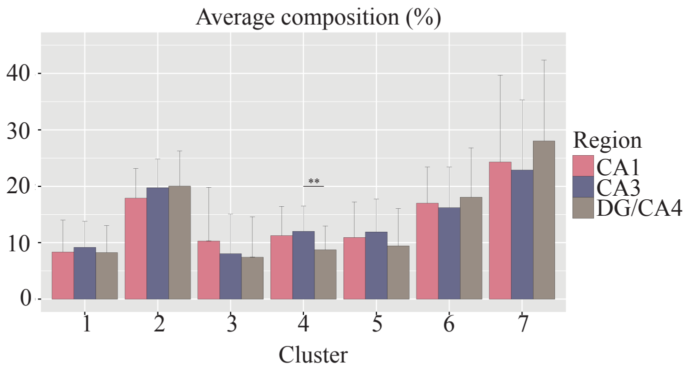
Average composition (%)
40
30
20
10
0
**
Region
CA1
CA3
DG/CA4
1
2
3
4
5
6
7
Cluster

Supp Fig 8
